# Body movement perception is shaped by both generic and actor-specific models of human bodies

**DOI:** 10.1101/2025.02.15.636568

**Authors:** Antoine Vandenberghe, Gilles Vannuscorps

## Abstract

Body movement perception is shaped by knowledge of the human body biomechanics. Apparent motion elicited by rapidly alternating pictures follows the shortest path between two body postures only if it is biomechanically plausible. And although we tend to perceive the location of a moving body part as slightly shifted forward along its trajectory, this perceptual extrapolation is absent (or reduced) if the movement would have been unlikely to continue along the same trajectory because of the body biomechanical constraints. The received view is that perception is shaped by a model of the observer’s own body. Here, we report three lines of evidence suggesting otherwise. First, we report the typical influence of knowledge of the upper limb biomechanics on apparent movement perception and perceptual extrapolation in two individuals born completely without (upper and lower) limbs (Experiments 1 and 2). Second, we report that we failed to find evidence that observers’ own biomechanics influence perceptual extrapolation of body movements (Experiment 3). Third, we show that participants’ perception is influenced by their knowledge of actor-specific biomechanics (Experiments 4- 8). We conclude that body movement perception is shaped by visually acquired models of both generic and actor-specific body biomechanics.

**Public significance statement:** Although information about others’ body movements reaches our visual system with a considerable delay, we can interact with them in real-time. Although we often blink, we perceive uninterrupted body movements. To solve the problems posed by informational lag and intermittence, the visual system appears to rely on knowledge of the human body biomechanics to extrapolate the most likely future (or unseen) position of observed limb movements. For the last 20 years, the received view has been that perceptual “filling-in” and extrapolation of others’ body movements are underpinned by a model of the observer’s own body, acquired through motor experience and proprioceptive feedback. Here, we report three lines of evidence indicating that body movement perception is rather shaped by visually acquired models of both generic and actor-specific body biomechanics. These findings provide new insight into how the human visual system extrapolates future body positions.

---

Although information about others’ body movements reaches our visual system with a considerable delay, we can interact in real-time. Although we often blink, we perceive uninterrupted body movements. To solve the engineering problems posed by informational lag and intermittence, the visual system appears to rely on models of the human body biomechanics to extrapolate the most likely future (or unseen) position of observed limb movements. Accordingly, apparent motion of body movements – the perception of body motion from a sequence of body postures – typically follows a biologically plausible path (Shiffrar & Freyd, 1990, Figure 1A). And although we tend to perceive the location of a moving body part as slightly shifted forward along its trajectory, this is not (or less) the case if the movement would have been unlikely to continue along the same trajectory due to the biomechanical constraints of the body (Vannuscorps & Caramazza, 2016a; Vandenberghe & Vannuscorps, 2023; Figure 1B). An outstanding issue concerns the nature of these models.

**Figure 1.**
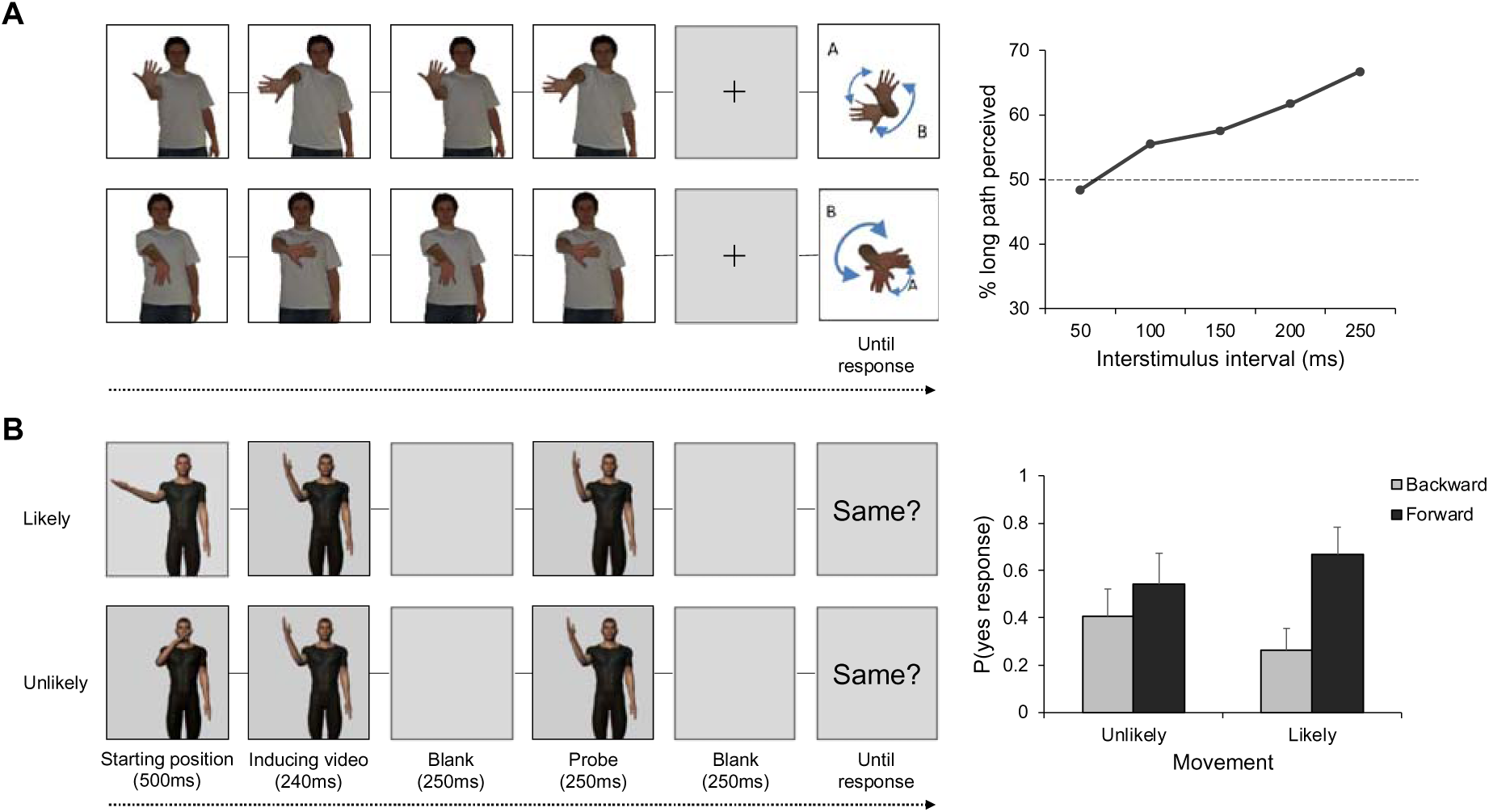
Illustration of the two main tasks that have revealed that perception of human body movements is shaped by knowledge of the human body biomechanics. *Note*. A. In the apparent motion task, participants see photographs of a human model whose body alternates between two postures at different frequencies and are asked to report the perceived path of movement, if any. If body movements were processed like any other object, the actor’s hand should be perceived as moving back and forth over the shortest path between these two postures (Shepard, 1981; Farrell & Shepard, 1981). In contrast to this hypothesis, when the shortest path between the postures is biomechanically impossible (as illustrated in the figure) participants often report perceiving a longer but biomechanically plausible path of motion (here, path B), and this bias increases with the duration of the interstimulus interval (results from Vannuscorps & Caramazza, 2016b). B. In the localization task, participants are asked to decide whether a photograph (probe) depicts an actor’s body in the same posture as it was at the end of a preceding video clip (inducing video). The actor’s posture in the photograph may be slightly shifted forward along the initial movement trajectory (“forward probe”) or slightly shifted backward (“backward probe”). When the movement displayed on the video clip would have been likely to continue along the same trajectory (likely condition), participants typically respond significantly more often “yes” for the forward than for the backward probes (i.e., they show “forward displacement” – also often referred to as “representational momentum”). However, this difference in the probability of responding “yes” to the two types of probes (forward – backward; “skewness index”) is typically smaller or absent when the movement would have been unlikely to continue along the same trajectory. This difference in skewness index for the two types of movements indicates an influence of knowledge of the body biomechanics on participants’ perception (results from Vannuscorps & Caramazza, 2016a).

For the last 20 years, the received view has been that perceptual “filling-in” and extrapolation of others’ body movements are underpinned by an internal model of the observer’s own body, originally acquired through motor experience and proprioceptive feedback to support the planning and control of movements (e.g., Knoblich et al., 2002; Wilson & Knoblich, 2005). In turn, the finding that perception of others’ bodies and movements is shaped by several rules and constraints that also affect the control and execution of actual movements is itself widely considered compelling evidence for this “motor hypothesis” of body movement perception (Grush, 2004; Hommel et al., 2001; Knoblich, 2008; Stevens et al., 2000; Wilson & Knoblich, 2005).

An alternative, however, is that visual perception is constrained by models of others’ bodies, learned through visuo-perceptual experience (Tessari et al., 2010; Vannuscorps et al., 2012; Vannuscorps & Caramazza, 2016a). Here we report three lines of evidence in support of this “visual hypothesis”. First, we report a typical influence of knowledge of the upper limb biomechanics on apparent movement perception (Experiment 1) and perceptual extrapolation (Experiment 2) in two individuals born without upper and lower limbs, who therefore may only have acquired this knowledge through the visual experience of viewing others. Second, we report a failure to find evidence that typically developed participants’ body abilities influence their perception (Experiment 3). Third, we show that typically developed participants’ perception of others’ body movements is shaped by actor-specific biomechanical knowledge (Experiments 4- 8).

## Transparency and openness

This work was not preregistered. The hypotheses, predictions, sample size, procedure, stimuli, exclusion criteria, and analysis pipeline of all the experiments were defined a priori. The stimuli and analysis pipeline were identical to those used in our previous studies (Experiment 1: Vannuscorps & Caramazza, 2016b; Experiment 2: Vannuscorps & Caramazza, 2016a; Experiments 3-8: Vandenberghe & Vannuscorps, 2023). Below, we detail how sample sizes were determined and provide information about the number of participants who were excluded from the sample based on a priori exclusion criteria in the method and analysis sections of all the experiments. Data were analyzed using SPSS 25 (IBM Inc.). All the stimuli and data have been made available on OSF (https://osf.io/yhp94/?view_only=12377ba3023e43508fe9926395c16af2).

All the experiments reported below were approved by the biomedical Ethics Committee of the Cliniques universitaires Saint-Luc, Brussels, Belgium (#B403201835404), and all participants gave written informed consent before experimentation.

## Experiment 1

If the influence of knowledge of the body biomechanics on perception reflects the recruitment of the observers’ own somatosensory and motor representations, then, knowledge of the body biomechanics should not influence the perception of upper limb movements in individuals congenitally deprived of upper limbs. In contrast, if this influence reflects internal models of others’ bodies acquired through visual experience, these individuals’ response profiles should be analogous to those of typically developed participants. To set apart these conflicting predictions, we tested the perception of apparent upper limb movements (Figure 1A) in two individuals born without upper and lower limbs and compared their perception to that of typically developed participants.

### Method

#### Participants

We tested two individuals born with severely shortened or completely absent upper and lower limbs (individuals with quadruple amelia, IQAs). IQA1 is a 60-year-old woman. She has a bachelor’s degree. She has a congenital quadruple amelia due to in-utero thalidomide exposure. On both sides, her upper limbs are constituted of two fingers positioned a few centimeters below the shoulders (no hands, arms or forearms are present). These fingers cannot be bent and can only be moved a few centimeters away or toward each other, allowing a “squeezing” movement. Both lower limbs are constituted of a short segment (left: ± 25 cm; right: ± 15 cm) followed by a foot composed of extra numerous toes (left: seven; right: six). There is no knee or ankle and only the left hip joint is functional. The foot and “leg-like” segment cannot be moved voluntarily but the toes appear to have typical mobility. Physical and mental development was otherwise normal. IQA1 reports that she used prosthetic legs from 3 to 14 years old and cosmetic legs until about 26 years old. She also used gas-powered upper limb prostheses between 3 and 7 years old, allowing her to act on split hooks through cables and pulleys. There is no reported history of phantom limb sensations or movements. She has a slight presbyopia that may be corrected by glasses. However, she is not able to put on her glasses herself and was not able to use them during the experiments. IQA2 is a 22-year-old man with a high-school degree, who has a history of learning disorders (especially reading and arithmetic). He has a congenital quadruple amelia of unknown origin characterized by a complete absence of upper and lower limbs. There is no history of prosthetic use but a lifelong history of very vivid phantom sensations of the four missing limbs. Although there is no history of serious neurological disorder IDA2 reports having experienced numerous concussions. He has a slightly corrected myopia.

IQA1 and IQA2’s responses were compared to that of 15 typically developed control participants (all right-handed, 13 females and two males; *M*_age_ = 22.8; range = 20-28) whose profile in this experiment was reported in a previous study (Vannuscorps & Caramazza, 2016a).

#### Material and procedure

The experiment was controlled by the online testable.org interface (http://www.testable.org), which allows precise spatiotemporal control of online experiments. IQA1 was tested on the 1536×864 pixels screen of her computer. IQA2 was tested on the 1080×1920 pixels screen of his smartphone. IQA1 and 2 were tested remotely under the supervision of the last author (GV) through a Visio conference system. During the experiments, IQA1 sat comfortably at about 60 cm from the screen while IQA2 laid on the floor next to his smartphone. At the beginning of each experiment, the participant was instructed to set the browsing window of the computer/tablet to full screen, minimize possible distractions (e.g., TV, phone, etc.), and position themselves as comfortably as possible in front of the screen.

Stimuli consisted of four pairs of photographs depicting an actor who remained stationary except for the position of his right (two pairs of photographs) or left (two pairs of photographs) upper limb (Figure 1A). In the four pairs of stimuli, the shortest path between the two hand positions was biomechanically impossible.

During the experiment, participants were first exposed to the rabbit-duck picture (Brugger & Brugger, 1993) and told “this stimulus can be perceived as a duck or as a rabbit. If a person was shown 100 times this same picture, it is likely that s/he would sometimes perceive the rabbit and sometimes the duck. Obviously, none of these responses would be correct or incorrect. This phenomenon demonstrates that the perception of the same stimulus can differ from time to time and from person to person. In the experiment that will follow, we are interested in the kind of movements that people perceive when they see rapidly presented pictures of a human body in different positions. This is also an example of perceptual ambiguity. Sometimes you may see a movement going in one direction, sometimes in another direction. There is no correct or incorrect response. We simply ask you to relax and tell us what you see.” This phase was added after pre-tests to avoid biasing participants from thinking about the plausibility of the perceived movements presented in the experiment (Vannuscorps & Caramazza, 2016b).

Subjects then performed a block of eight maximal amplitude trials followed by a block of 80 trials (2 sides x 2 movements x 5 ISIs x 4 iterations) in which stimuli were pseudo-randomly mixed. Each trial started with the presentation of a blank screen for 500 ms followed by four cycles of two different hand positions separated by one blank screen. These cycles were presented at five different presentation speeds. The fastest speed involved presenting the photographs for 100 ms separated by a 50 ms interstimulus blank screen. The second, third, fourth, and fifth presentation rates involved presenting the photographs for 150, 200, 250, and 300 ms and the blank screen for 100, 150, 200, and 250 ms, respectively. After the last photograph of each trial, the participants viewed a blank screen for one second and then a response figure showing two possible paths of apparent motion between the actor’s hand positions (e.g., see Figure 1A). This figure was displayed until a response was provided. There were two versions of each response figure. In one version, the shortest path was labeled “A” and the longest “B”. In the other version, the assignment of letters was reversed. When the response figure appeared, participants were asked verbally whether they had perceived a movement, and if they had, whether the perceived movement corresponded to one of the two movements displayed on the figure, and indicate which one by naming the corresponding letter, or whether they had perceived some other movement, and to describe it. The response was recorded by the experimenter. The encoding of the response launched the next trial. This experiment may be visualized at http://www.testable.org/experiment/7/919310/start.

### Results

The results of all participants are displayed in Figure 2. We conducted two series of analyses. We first looked for the presence of the typical bias toward the perception of a long and biomechanically plausible path modulated by the ISI in the response profiles of IQA1 and IQA2. To do so, we tallied the number of trials in which IQA1 and IQA2 reported perceiving the long path at each ISI. As shown in Figure 2, both IQA1 and IQA2 often reported perceiving the long (but biomechanically plausible) path of motion (IQA1: 76%; IQA2: 55%), and their percentage of perceived “long paths” increased with the ISI. This increase was statistically significant as they both reported a significantly larger number of perceived “long paths” at the longest (300 ms) than at the shortest (100 ms) ISI (both χ*²* (1) > 5.23, both *p*s ≤ .02).

**Figure 2.**
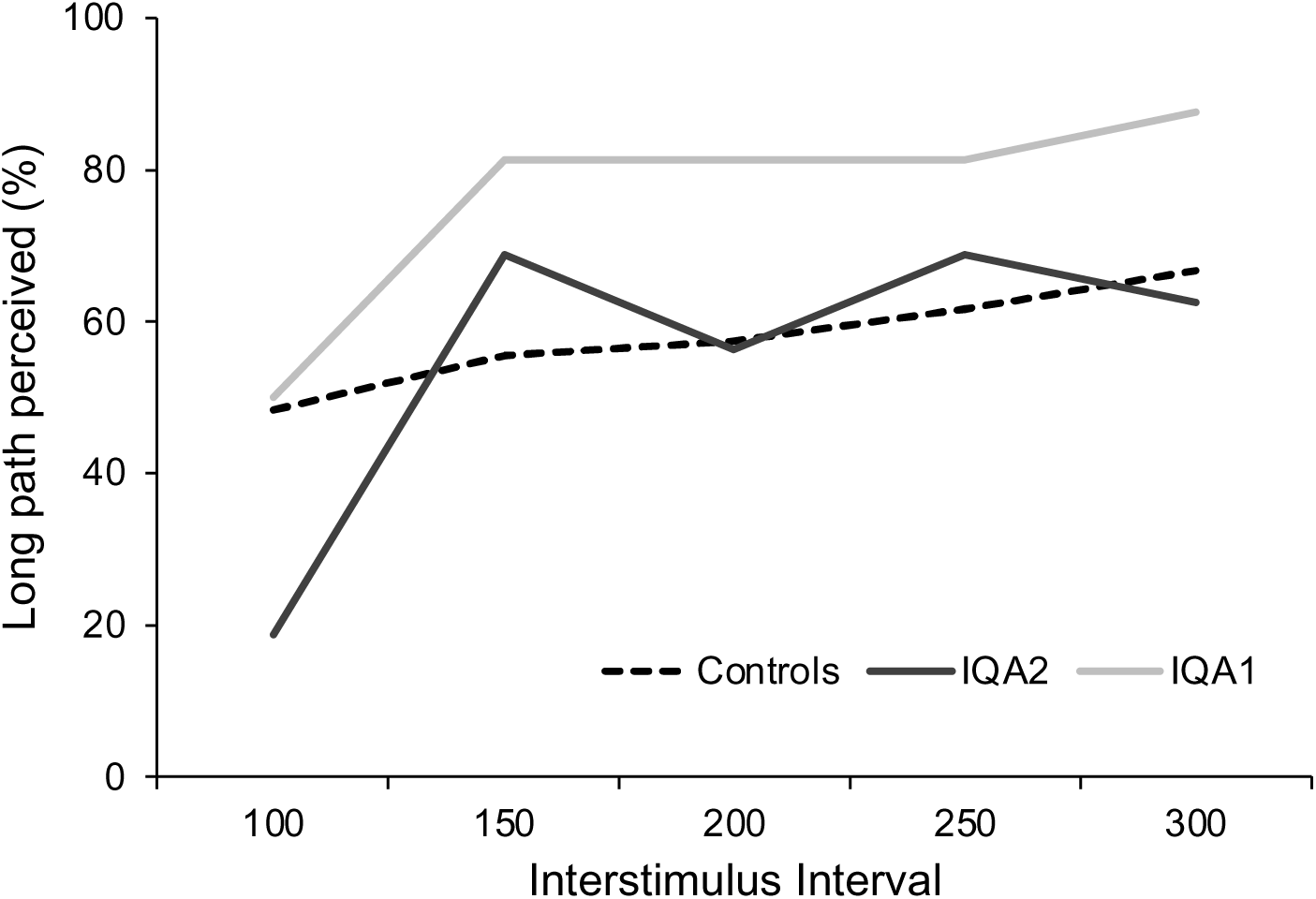
Results of Experiment 1. *Note*. The Y-axis represents the percentage of “long paths” chosen by IQA1, IQA2, and control participants at each level of the interstimulus interval.

Then, we compared their performance to that of the control participants to investigate the possibility that their performance nevertheless differed from that of typically developed participants on these two indexes. In contrast to this possibility, IQA1 and IQA2’s tendency to show a bias toward biomechanically possible movement did not differ from that of controls (modified *t*-tests, IQA1 *t*(14) = 0.70, *p* = .49, and IQA2 *t*(14) = −0.12, *p* = .90). The increase of the bias between the shorter and the longer ISI did not differ from that of controls (modified *t*- tests, IQA1 *t*(14) = 1.04, *p* = .32, and IQA2 *t*(14) = 1.37, *p* = .19).

### Discussion

IQA1 and IQA2’s responses in Experiments 1 were characterized by the two typical indexes of an influence of knowledge of the upper limbs’ biomechanical constraints on apparent motion perception. First, they reported perceiving the actor’s hand moving along the long and biomechanically plausible path as often as the controls. If they perceived the upper limb movements like any other objects (Farrell & Shepard, 1981; Shepard, 1981), or if the shortest path was biomechanically possible (Shiffrar & Freyd. 1990, 1993), they would have consistently perceived the actor’s hand as moving back and forth over the shortest path between these two postures. Second, like the controls, their perception of the longer path increased with the ISI (Shiffrar & Freyd, 1990, 1993). Increasing the ISI results in the perception of longer paths only when the shortest path is biomechanically impossible (Shiffrar & Freyd, 1990, 1993). Hence, this demonstrates that the responses of IQA1, IQA2 and of the controls were not generated at random.

This finding corroborates those of previous studies that showed that individuals born without upper limbs may show a typical biomechanical bias in the apparent motion (Funk et al., 2005; Vannuscorps & Caramazza, 2016b). The present finding goes beyond this previous evidence, however, by demonstrating these effects in individuals deprived of both upper and lower limbs. In individuals born without upper limbs, the observation of hand actions recruits part of the motor system naturally used to execute lower limb actions (Aziz-Zadeh et al., 2012; Gazzola et al., 2007; Vannuscorps et al., 2019). Although hand and foot movements are constrained by different skeletal and muscular features, they have some biomechanical similarities, such as a smaller degree of freedom of rotational movements away than toward the body midline. Hence, it could be that these previous individuals’ perception of upper-limb movements was somewhat constrained by an internal model of their lower limbs. The present results are immune to this limitation and demonstrate that the effect of knowledge of the body biomechanics on body movement perception does not require motor experience *at all*, that is, neither with the observed body part nor with another body part that may serve as an analogue. This constitutes existence proof that the influence of knowledge of the body biomechanics on body movement perception can rely exclusively on models developed through visual experience.

## Experiment 2

Experiment 2 aimed at validating the findings of Experiment 1 through conceptual replication. We used a localization task to test both control participants and IQA1 and IQA2’s perception of a body movement that would be easy to continue along the same trajectory beyond its endpoint (the “likely” movement in Figure 1B), and of a body movement that would be difficult to continue along the same trajectory beyond its endpoint by an average actor (the “unlikely’ movement in Figure 1B). In localization tasks, the influence of knowledge of the body biomechanics on perception is indexed by a difference between the skewness index (probability of yes response for the forward probes minus that for the backward probes) associated with these two types of movements (see caption of Figure 1 for explanation). Typically, this difference emerges because the perception of a “likely” movement evokes forward displacement (i.e., a positive skewness index significantly different from zero) while the perception of an “unlikely” movement does not, or significantly less. Thus, if the influence of knowledge of the body biomechanics in localization tasks reflects the recruitment of the observers’ own somatosensory and motor representations, then, IQA1 and IQA2’s response profiles should be similar for the two movements (i.e., there should be no difference in skewness index between the two movements). In contrast, if this bias reflects an influence of models of others’ bodies, acquired through visual experience, then IQA1 and IQA2’s response profiles should be analogous to those of typically developed participants.

### Method

#### Participants

IQA1 and IQA2’s responses were compared to that of 32 typically developed and able control participants (26 females, six males, *M_age_*= 21, *SD* = 3.8), a sample size providing a power of 95% to detect an effect of a size of Cohen’s d of .6 – the effect size found in previous studies (Vannuscorps & Caramazza, 2016a; Wilson et al., 2010). All of them received course credit for their participation.

#### Material and Procedure

Participants performed the experiment in the same conditions as in Experiment 1. Stimuli consisted of two video clips computed from frames originally used in Vannuscorps and Caramazza (2016a; Experiment 7; cf. Figure 1B). They depicted a computer-generated actor facing the participant, with the right arm raised to form a 60° angle with the body and flexed to form a 90° angle between the arm and the forearm. In the first video (likely condition on Figure 1B), a first frame depicting the actor’s right forearm tilted 80° counterclockwise from the midsagittal line was followed by a 60° internal rotation of the right shoulder (i.e., toward the body midline). In the second video (unlikely condition on Figure 1B), a first frame depicting the actors’ forearm tilted 40° clockwise from the midsagittal line was followed by a 60° external rotation of the right shoulder (i.e., away from the body midline). The actor’s body, left arm, and face remained still. The last frame of the two video clips was identical and depicted the hand positioned at the average limit of the amplitude of external rotation of the shoulder, that is, 20° away from the midsagittal line (Gill et al., 2020; Hill et al., 2010). The first frame of the video was presented for 1,000 ms and then, the movement was depicted by presenting 12 frames on which the rotation of the forearm was increased by steps of 5° for 33 ms each (150 °/s; 400 ms in total). This way, the likely video clip depicted a smooth and continuous rotating movement of the arm of constant velocity that would be likely to continue along the same trajectory after the position of the hand on the last frame. However, the unlikely video clip depicted a rotating movement of the arm of constant velocity that would be unlikely to continue along the same trajectory after the position the hand reached on the last frame, due to the biomechanical constraints imposed by the joints of the shoulder. The set of stimuli also included three pictures, used as probes in the experiment. The first picture was the last frame of the video clips. This picture will be referred to hereafter as the “identical” probe. The two other pictures depicted the hand of the actor slightly (4°) shifted away from or toward the actor. This shift (4°) was selected after pilot experiments because it maximized the number of errors for the forward probes while minimizing the number of errors for the backward probes (i.e., it maximized the task sensitivity). Each of these pictures will be referred to hereafter as either the forward or the backward probe depending on whether it depicts the hand of the actor shifted slightly forward or backward along its initial trajectory.

Participants performed three blocks of 60 trials. In each block, the two conditions and the three different probes were in equal proportion and mixed in a pseudo-randomized order. The first block included 12 practice trials with the three probes repeated evenly. Within each block, each trial started with the presentation of the video clip, a 250 ms blank screen, one of the three probe pictures displayed during 250 ms and, finally, the phrase “Was the hand on the picture exactly at the same place as it was at the end of the video-clips?” and, below it, “If yes, say yes; If no, say no” for patients, and “If yes, press yes; If no, press no” for control participants until a response was recorded.

### Results

As shown in Figure 3, IQA1, IQA2 and most control participants showed a positive skewness index in the LIKELY movement condition, but this effect was either smaller (in IQA2) or reversed (in IQA1) in the UNLIKELY movement condition.

**Figure 3.**
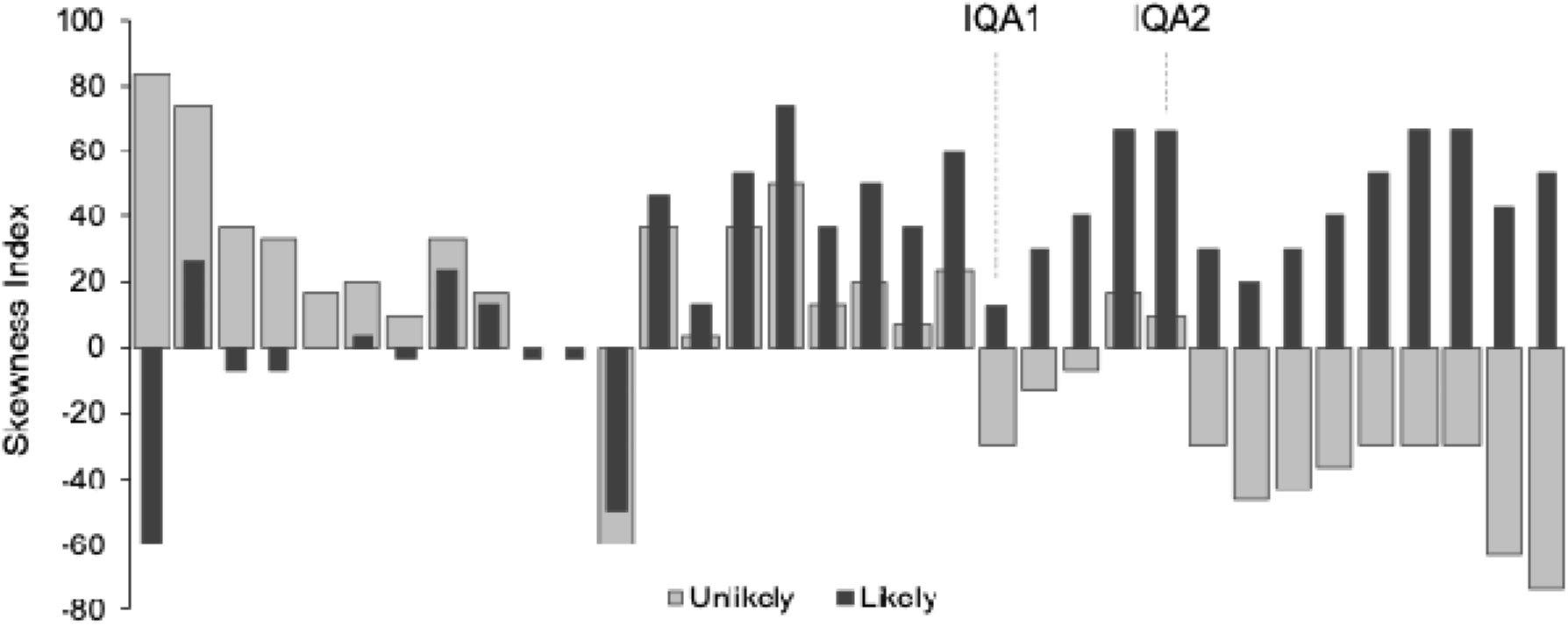
Results of Experiment 2. *Note*. Individual patterns of responses in Experiment 2. The Y-axis represents the Skewness index of each participant, computed by subtracting the percentage of yes responses for backward probes from that for forward probes, as a function of movement conditions. Responses of IQAs are labeled in the graph. Participants are arranged in ascending order based on the disparity between the magnitude of forward displacement observed in the likely and unlikely conditions. Individuals on the rightmost side are those whose responses show the largest influence of the knowledge of upper-limb biomechanics.

We first tested whether this apparent difference in skewness index was statistically significant in the control participants. To do so, their responses were inserted in a generalized linear mixed model (GLMM) with participants’ responses (yes/no) as a dependent variable, and movement (likely/unlikely) and probe (forward/backward) as fixed within-subject factors. Participant was included as a random factor. As expected, we found a significant interaction between movement and probe, *F*(1,3836) = 65.95, *p* < .001, *OR* = 3.13.

Then, to characterize more precisely the performance of the control participants, we conducted additional analyses to explore whether participants’ perception was characterized by forward displacement (i.e., a significantly larger number of “yes” responses for forward than for backward probes). This was the case for the movement that would have been likely to continue (*M* = .29, *SE* = .02), *t*(3836) = 12.38, *p* < .001, but not for the movement that would have been unlikely to continue (M = .02, SE = .02), *t*(3836) = 0.92, *p* = .34.

Because we used only one forward and one backward probe, in theory, the smaller skewness index and non-significant forward displacement associated with the unlikely movement could either reflect a smaller or a larger forward displacement, peaking beyond the position depicted on our forward probe.

To tell apart these two interpretations, we compared the probability of “yes” responses for the backward probes in the two movement conditions. The unlikely movement was associated with a significantly larger probability of “yes” responses for the backward probes than the likely movement, *t*(3836) = 8.45, *p* < .001. The result is incompatible with the hypothesis that the unlikely may in fact evoke a significantly larger forward displacement than the likely condition.

To test whether IQA1 and IQA2’s perception was also influenced by knowledge of the body biomechanical constraints, we performed binomial logistic regression analyses on each participants’ responses (yes/no) as the dependent variable and movement condition (likely/unlikely) and probe (forward/backward) as independent variables. In line with this possibility, the results of these analyses indicated a significant interaction between these two independent variables in both participants, both χ² (1) > 4.92, both *p*s < .02, both *OR*s > 1.41. Additional analyses conducted to characterize the performance of IQA1 and IQA2 in this task further indicated that this interaction emerged in the context of a significant forward displacement in the likely movement condition in IQA2 (χ² (1) = 38.2, *p* < .001; but not in IQA1: χ² (1) = 1.15, *p* = 0.28) and of a significant backward displacement in the unlikely movement condition in IQA1 (χ² (1) = 6.08, *p* = .01; but not in IQA2: χ² (1) = 2.09, *p* = 0.15).

A series of additional analyses further indicated that the biases found in IQA1 and IQA2 were similar to those found in control participants. A series of modified *t*-tests (Crawford & Howell, 1998) showed that the size of the displacement in the likely condition (modified *t*-tests, IQA1 *t*(31) = −0.41, *p* = .68, and IQA2 *t*(31) = 1.21, *p* = .23) and the size of the displacement in the unlikely condition (modified *t*-tests, IQA1 *t*(31) = -0.82, *p* = .41, and IQA2 *t*(31) = 0.20, *p* = .84) did not differ significantly between the controls and in the two IQAs. Bayesian Standardized Difference tests (Crawford & Garthwaite, 2007) further indicated that the difference between the two conditions was similar in the IQAs and in the controls (BSDTs, IQA1 *p* = .75 and IQA2 *p* = .52).

Note, finally, that we ran additional exploratory analyses of the data reported in this section, including analyses of variances (ANOVAs) and GLMMs that include the independent variables and their interaction as random factors. The results of these additional analyses, which may be found on OSF, led to the same conclusions.

### Discussion

If the influence of knowledge of the body biomechanics in localization tasks reflected the recruitment of the observers’ own somatosensory and motor representations, then, IQA1 and IQA2’s response profiles should have been similar for the two movements (i.e., there should be no difference in skewness index between the likely and the unlikely movements). At odds with this prediction, IQA1 and IQA2’s response profiles for the two types of movements differed significantly. Like the controls, IQA1 and IQA2 both showed a significantly larger skewness index for the movements that would have been likely than for the movements that would have been unlikely to continue along the same trajectory. In IQA1, this difference emerged in the context of a significantly larger number of “yes” responses to backward than forward probes for the movement unlikely to continue beyond its endpoint, and in IQA2, in the context of a significantly larger number of “yes” responses to forward than backward probes for the movement that was likely to continue beyond its endpoint. These results support the visual hypothesis according to which visual perception is constrained by models of others’ bodies, learned through visuo-perceptual experience.

In addition to predicting that the two movements would evoke different percepts in the controls and in IQA1 and IQA2, the visual hypothesis also suggests that this perceptual difference should manifest as a larger forward displacement in the likely movement condition than in the unlikely one. The results supported this prediction. First, in both the controls and in IQA2, observing the likely movement led to a significantly higher number of “yes” responses to the forward probes than to the backward probes, indicating forward displacement. Indeed, it is unclear how this finding could emerge if the observation of the likely movement did not elicit forward displacement, or if it is associated with backward displacement.

Second, and more importantly, the perception of the likely movement evoked a significantly larger forward displacement than the perception of the unlikely movement. This conclusion follows from the fact that the perception of the likely movements was associated with (1) forward displacement (see above), (2) a larger skewness index and, (3) a significantly smaller probability of “yes” responses to the backward probes. Again, it is unclear how this combination of effects could be accounted for by any other hypothesis (i.e., if the perception of the likely movement was not associated with a larger forward displacement or if it was associated with a smaller forward displacement).

A larger displacement for movements directly away than towards the body could also result from “landmark attraction”, i.e., a tendency to judge the distance from a target to a landmark (here, the body of the actor) as being smaller than the distance from the landmark to that target (Bryant & Subbiah, 1994) that has been shown to reduce the size of displacement when an object moves away from a landmark than toward it (Hubbard & Ruppel, 1999). However, the effect of landmark attraction on displacement for objects moving away versus toward a landmark has been shown to be significantly smaller than the effect of knowledge of the body biomechanical constraints on displacement for body parts moving away versus toward the body (Vandenberghe & Vannuscorps, 2023). Hence, although landmark attraction is likely to contribute to the difference in skewness index for the two types of movements found in IQA1 and IQA2 (and in the controls), it cannot easily account for the fact that this difference was of a similar size (in fact slightly above average) than the difference found in the control participants.

This finding corroborates those of previous studies that showed that individuals born without upper limbs may show a typical biomechanical bias in localization tasks (Vannuscorps & Caramazza, 2016a). However, our results extend this evidence by demonstrating these effects in individuals lacking both upper and lower limbs, suggesting that these effects may emerge without any motor experience with either the relevant body part (the hand) or one that may serve as an analogue (the foot).

Yet, it could be that visual models of others’ bodies are built only in individuals who cannot engage their own internal body models. To address this issue, in Experiments 3-7 we tested additional conflicting predictions of the motor and visual hypotheses in typically developed participants.

## Experiment 3

Experiment 3 aimed at testing two differential predictions of the motor and visual hypotheses. The first concerned the origin of the biomechanical bias previously documented in localization tasks (Experiment 2; Wilson et al., 2010; Vandenberghe & Vannuscorps, 2023). If this biomechanical bias reflects the recruitment of the observers’ own somatosensory and motor representations, then, forward displacement associated with the unlikely movement should be larger in observers who would be able to continue this movement beyond its endpoint than for observers who would not be able to do so. As a consequence, the discrepancy between the skewness index associated with the two movements depicted in Experiment 2 (see also Figure 1B) should be larger in participants who would be able to continue only one of the two movements along the same trajectory than in individuals who would be able to continue both movements. In contrast, if this bias reflects knowledge or hypotheses about how other people are likely to move, the discrepancy in skewness index between the two movements should not be influenced by the observer’s own ability.

The second concerns perceptual extrapolation of body movements. Typically, the biomechanical bias emerges in localization tasks because the perception of movements that would be easy to continue along the same trajectory induces forward displacement (i.e., significantly larger number of “yes” responses for the forward than for the backward probes) while this is not, or less, the case for movements that would be difficult to follow along the same trajectory. If perceptual extrapolation (forward displacement) depends on the observers’ own somatosensory and motor representations, then it should be larger when a movement is viewed by an observer who would be able to continue that movement along the same trajectory, than when it is viewed by the observer who would not.

### Method

#### Participants

We used a convenience sample of 74 participants (57 females, 17 males, *M_age_* = 21; *SD* = 1.8) whose performance in the localization task was measured in a previous study (Vandenberghe & Vannuscorps, 2023). Although their body abilities were measured in the context of their participation in that previous study, this information was neither reported nor exploited before. All of them received course credit for their participation.

#### Material and procedure

Participants performed two tasks. First, they participated in a localization task that was very similar to that reported in Experiment 2. The stimuli and the procedure differed from those of Experiment 2 in three aspects: (1) we added a probe picture depicting the actor’s hand aligned with the midsagittal line and used it as catch trials in the “unlikely” condition (it corresponded to a 20 degrees backward displacement of the hand in this condition); (2) participants responded by pressing the keys “o” for yes or “n” for no (corresponding to the first letter of the equivalent words “oui” and “non” in French) and (3) there were 15 repetitions per probe (forward, identical, backward) and movement (internal vs. external rotation of the shoulder), and 15 catch trials in the external rotation (“unlikely”) condition, for a total of 105 trials. At the end, the participants were invited to guess the goal of the task. This task was conducted on Psychopy 1.90.3 (Peirce, 2007) and presented on a 60 Hz monitor.

In a second task, we measured participants’ maximal amplitude of external rotation of the shoulder with the body and arm positioned in the same posture as that shown in the videos of the first task (see Figure 1B). Participants sat on a chair in front of a table, with their right arm uncovered. The height of the chair was set such that when participants placed their right elbow on the table, their arm was rotated approximately 20° away from the sagittal plane and formed an angle of 60 degrees with the coronal plane. At the beginning of the trial, the palm of their right hand was positioned against a vertical platform aligned with the sagittal plane. As depicted in Figure 4, one leg of a digital goniometer was glued to that platform, while the second leg remained free to move. An angle bracket was attached to this second leg so that participants executing an external rotation of the shoulder would cause the second leg of the goniometer to rotate. Participants were asked to execute an external rotation of their right shoulder of maximal amplitude. During the execution of the movement, the experimenter made sure that the participant’s body and arm remained still while moving the right forearm.

**Figure 4.**
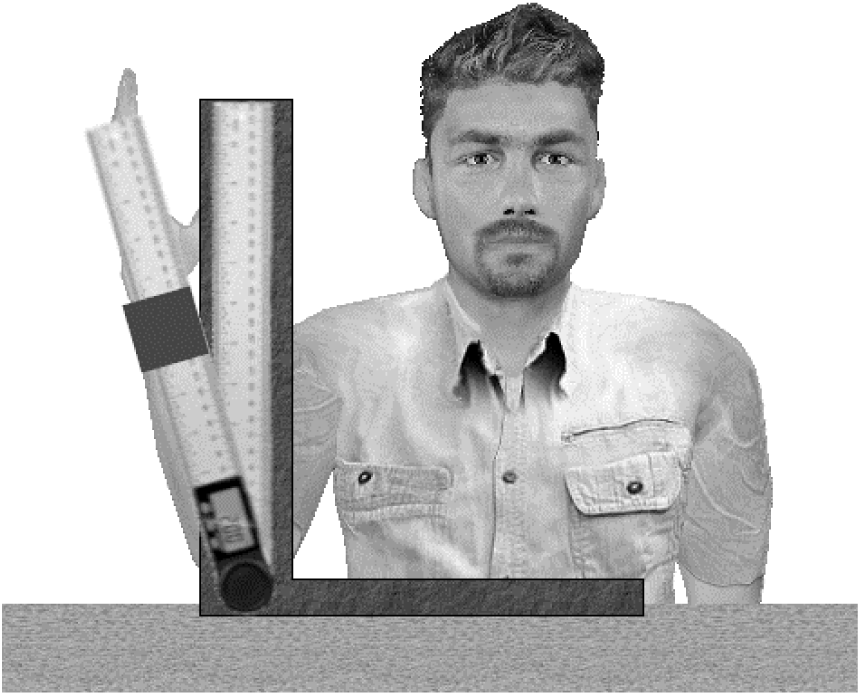
Maximal amplitude measurement setup. *Note*. The grey square represents the angle bracket attached to the digital goniometer.

### Analyses and results

Two exclusion criteria were predetermined: (1) failing to reject more than one catch trial or (2) correctly guessing the purpose of the experiment. No participant was excluded from the analyses.

To test whether the skewness index induced by the observation of a body movement is influenced by whether an observer would have been able or not to continue this movement along the same trajectory beyond the end-point depicted in the video, we first divided participants into two groups depending on whether their maximal amplitude of external rotation was below (37 participants, *M* = 14.8, *SD* = 3.3) or beyond (37 participants, *M* = 26.5, *SD* = 5.9) that shown in the video-clips (rigid/flexible groups).

Then, we ran a generalized linear mixed model (GLMM) with participants’ responses (yes/no) as a dependent variable, movement (internal/external) and probe (forward/backward) as a fixed within-subject factor, Group (rigid/flexible) as a fixed between-subject factor and Participant as a random factor to test whether the influence of biomechanical knowledge on perceptual extrapolation depends on the observers’ ability. At odds with the prediction of the motor hypothesis, we found a significantly larger skewness index in the internal condition than the external condition (i.e., a biomechanical bias) for both the rigid group, *F*(1,2216) = 76.64, *p* < .001, *OR* = 6.44, and the flexible group, *F*(1,2216) = 50.81, *p* < .001, *OR* = 4.30, and the interaction between Probe and Movement did not interact with the Group, *F*(1,4432) = 1.83, *p* = .18, *OR* = 1.50 (cf. Figure 5). Hence, we failed to detect a significantly larger biomechanical bias in participants who would be able to continue only one of the two movements along the same trajectory (rigid) than in individuals who would be able to continue both movements (flexible).

**Figure 5.**
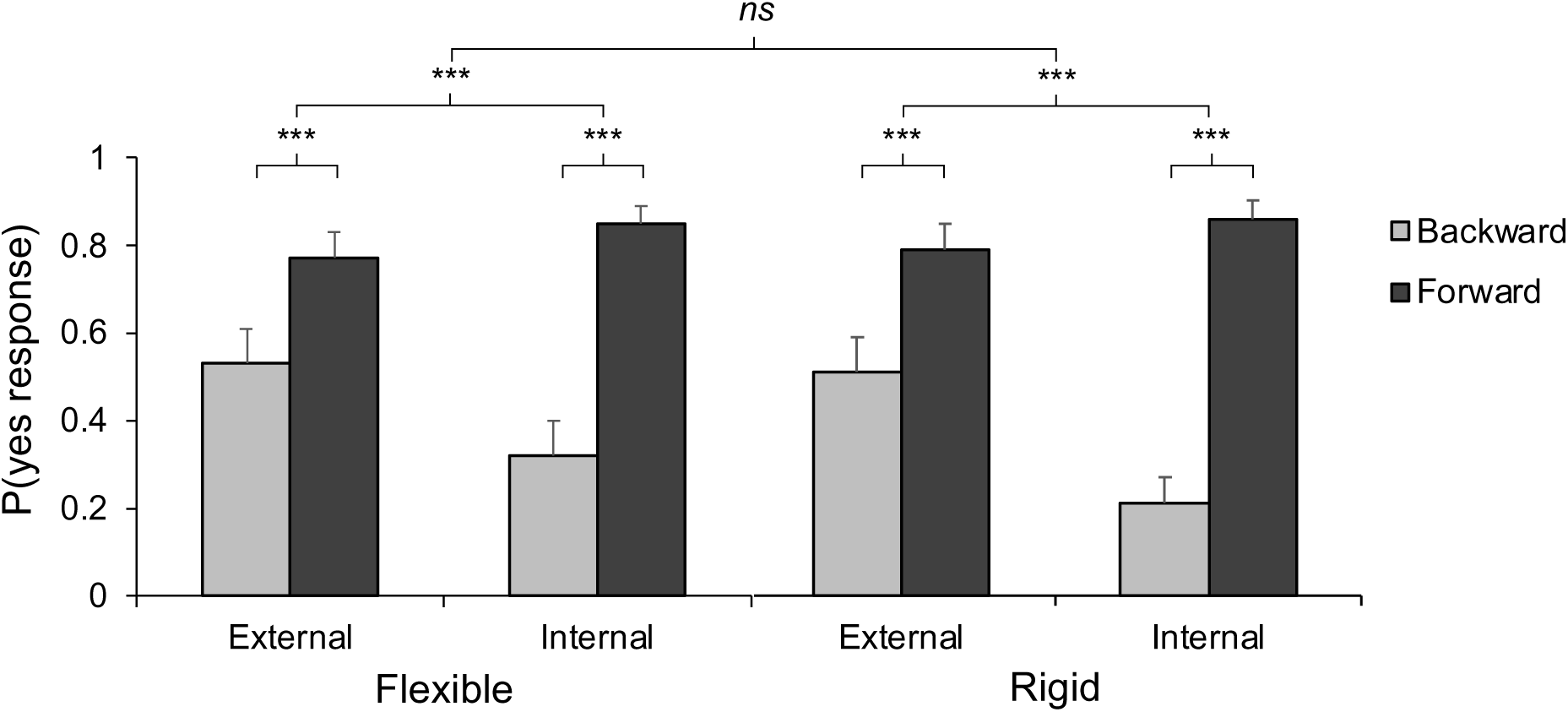
Results of Experiment 3. *Note*. Probability of a “yes” response as a function of Probe (forward/ backward), Movement (external/internal), and Group (flexible/rigid). Error bars depict 2 standard errors of the mean. *ns*: non-significant, \*\*\**p* < .001

Additional analyses performed on the data of the 20 most extreme participants from both groups led to the same conclusion (the results are available on OSF). Although the rigid and flexible groups had a maximal amplitude of movement far below and above (12 and 27 degrees, respectively) that shown on the external rotation stimulus (20 degrees), their average skewness index was identical (28% more “yes” responses for the forward than for the backward probes).

Then, we tested whether perceptual extrapolation depends on the observers’ own somatosensory and motor representations. A prerequisite was to verify that participants’ perception was indeed characterized by forward displacement in this task (i.e., a significantly larger number of “yes” responses for forward than for backward probes). This was the case for both movements in both groups, all four *t*s(4432) > 7.55, *p*s < .001. Then, we tested whether forward displacement for the external rotation was indeed larger for the flexible than for the rigid group. As discussed in the result section of Experiment 2, this would predict a combination of a significantly larger skewness index and of a smaller probability of “yes” responses to the backward probes for the flexible than for the rigid group. In contrast to this prediction, and as shown in Figure 5, the two groups of participants had a very similar skewness index, *F*(1,2216) = 0.77, *p* = .38, *OR* = 0.84, and provided a very similar probability of “yes” responses to the backward probes of the external rotation, *t*(2216) = 0.34, *p* = .74. Hence, a generalized linear mixed model (GLMM) with participants’ responses on the external rotation movement (yes/no) as a dependent variable, probe (forward/backward) as a fixed within-subject factor, Group (flexible/rigid) as a fixed between-subject factor and Participant as a random factor indicated that there was no significant interaction between Probe and Group, *F*(1,2216) = 0.77, *p* = .38, *OR* = 1.19. And a Bayesian Repeated Measures ANOVA carried out on the same data indicated that a model including only Probe as a within-subjects factor and Group as a between-subjects factor (H0) was 3.8 times more likely than an alternative model including also the two-way interaction between Probe and Group predicted by the motor hypothesis (H1).

Note, finally, that we ran additional exploratory analyses of the data reported in this section, including analyses of variances (ANOVAs) and GLMMs that include the independent variables and their interaction as random factors. The results of these additional analyses, which may be found on OSF, led to the same conclusions.

### Discussion

In line with the predictions of the visual hypothesis, but in contrast to the predictions of the motor hypothesis, we found (1) no evidence that the discrepancy in skewness index between the two movements was smaller in the flexible than in the rigid group and (2) no evidence that the size of forward displacement evoked by the external rotation of the shoulder could be larger in the flexible than in the rigid group.

The results of Experiments 1 and 2 indicated that the typical motion extrapolation and biomechanical bias that characterize body movement perception do not require somatosensory or motor representations corresponding to the observed movement (Experiments 1 and 2). The results of Experiment 3 additionally suggest that an observer’s somatosensory or motor representations either do not influence motion extrapolation or that their influence is not easy to detect. Of course, these findings do not imply that an observer’s own ability does not influence or modulate the size of the biomechanical bias at all and/or in any context. Our sample size did not provide us with enough statistical power to detect small effect sizes with a high probability. A larger sample size could have allowed us to detect a subtle influence of the observer’s own ability. Some aspects of our experimental design or some of our methodological choices (e.g., use of catch trials in the unlikely condition, speed of movement, mode of response) may also be responsible for our failure to detect this influence. Future studies will be needed to explore this possibility. In the meanwhile, our findings at the very least underscore the need for a shift in the burden of proof concerning the role of representations of one’s own body capabilities in perception. The aim of Experiments 4-7 was to test another (positive) prediction of the visual hypothesis.

## Experiment 4

In this experiment, typically developed and able participants performed a localization task in which they saw two different computer-generated actors performing rotations of the shoulder with different amplitudes. As shown in Figure 6, there were two types of trials. In the familiarization trials (FTs) the two actors performed movements of different amplitudes: larger for the “flexible” and smaller for the “rigid” actor. In the trials of interest (TIs), the two actors performed the same movement: an external rotation of the shoulder up to a point located well within the range of movement performed by the flexible actor in the FTs, but outside the range of movement performed by the rigid actor.

**Figure 6.**
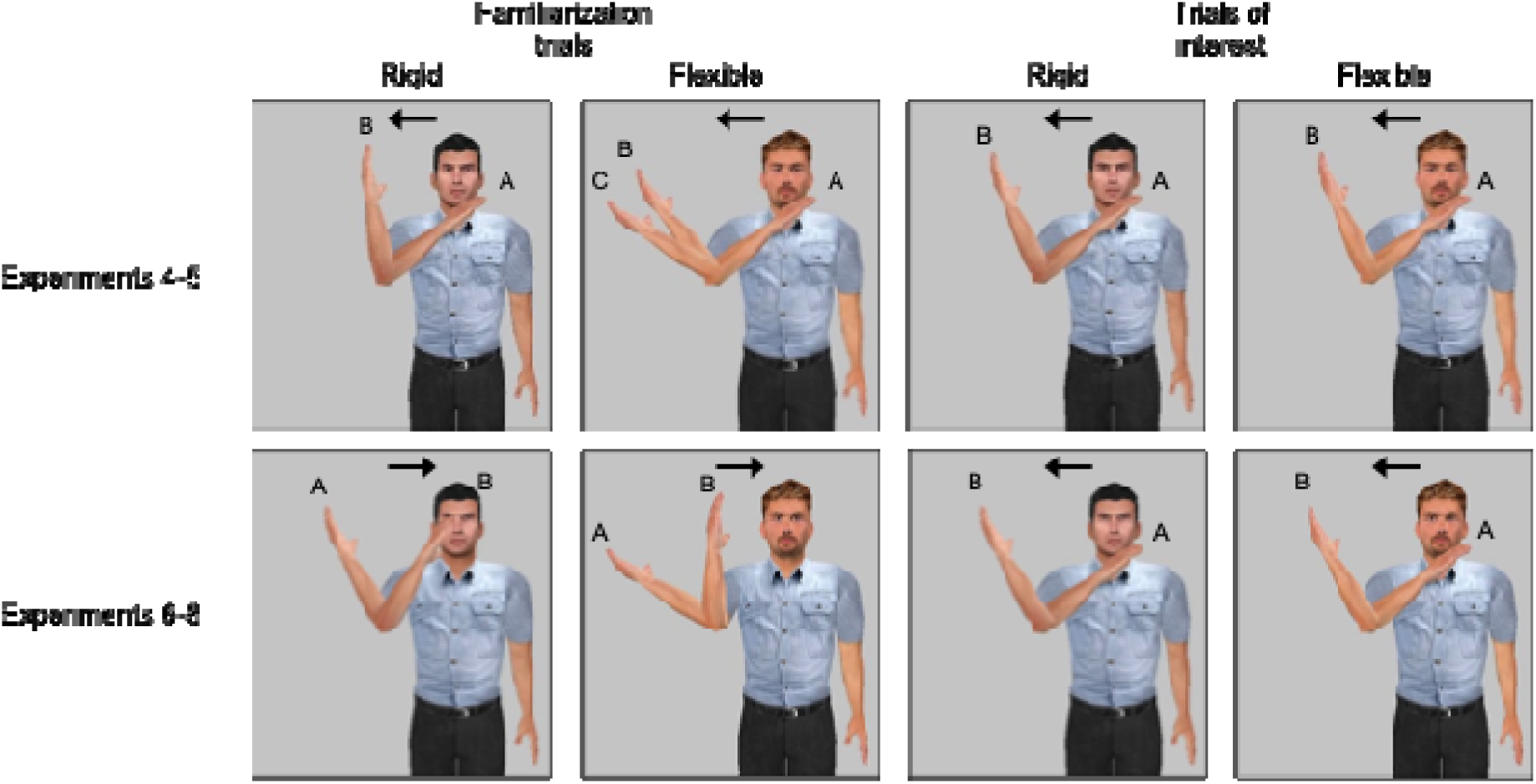
Types of trials by actor and experiment. *Note*. Starting (A) and final positions (B & C) for both actors in the familiarization trials (FTs) and trials of interest (TIs). The final position of the hand of the flexible actor in the FTs corresponded to positions B and C in Experiments 4 and 5, respectively.

The analysis of participants’ responses to the TIs allowed us to test a new differential prediction of the motor and visual hypotheses. According to the motor hypothesis, there should be no difference between the displacement evoked by the observation of the TIs executed by the two actors. If the visual hypothesis is correct, however, there should be a significant difference between the skewness index associated with the two actors. In particular, the perception of the TIs should evoke a larger forward displacement when it is executed by the flexible than when it is executed by the rigid actor.

### Method

#### Participants

A sample size of 20 participants (14 females, six males, *M_age_* = 23.1, *SD* = 3.6) was determined a priori on the basis of a power analysis. This provided a power of 82% to detect an effect of the actor of a size similar to that found in a pilot study involving 20 other participants (Cohen’s *d* = 0.6). All participants received course credit for their participation. All the participants received a compensation of seven euros for their participation.

#### Material and procedure

Stimuli depicted two different actors (the “black-haired” actor and the “blond-haired” actor, see Figure 6). They were divided into two sets (a familiarization and a test set). Each participant saw four video clips and 10 probe pictures. In each video clip, the first frame depicted the actor facing the participant with the right arm raised in the sagittal plane to form a 60° angle with the coronal plane, flexed to form a 90° angle between the arm and the forearm, and the right forearm tilted 50° clockwise from the mid-sagittal plane (cf. Figure 6). The two video clips of the familiarization set aimed to familiarize participants with each actor’s body ability. The “flexible” actor was shown performing a 90° external rotation of the right shoulder (i.e., away from the body midline), while the “rigid” actor performed only a 60° external rotation of the right shoulder. For half of the participants, the black-haired was the flexible actor and the blond-haired was the rigid actor, and vice-versa for the other half. The movement was depicted by presenting 18 frames (flexible actor) or 12 frames (rigid actor) on which the rotation of the forearm was increased by steps of 5° for 16.6 ms each (300 °/s; 300 ms and 200 ms durations). In the two video clips of the test set, both actors performed a 75° external rotation of the shoulder. These videos comprised 15 frames displayed for 16.6 ms each and lasted for 250 ms in total. The probes depicted either the final position of the actors on the familiarization video clips (identical probe, 2 pictures) or the hand of the actor displaced 5° forward or 5° backward from its position at the end of the four video clips (forward and backward conditions, 8 pictures).

During the experiment, we first showed to participants the last frame of the video clips of the familiarization set for both actors (the flexible and rigid actors were depicted with their right shoulder rotated by 90° and 60°, respectively) and told them explicitly that this corresponded to the two actors’ maximal range of motion. Accordingly, we explicitly referred to these two actors as “the flexible” and “the rigid” actors and told participants that this information “could help them” solve the task that they were to perform. Then, two tasks were explained to the participants. First, they were introduced to the localization task. Second, they were told that, in each trial of the localization task, the actor would initially remain still until they identified which actor was depicted on the frame – either the “flexible” (press the “f” key) or the “rigid” actor (press the “r” key). This second task was added to ensure that participants paid attention to the actor before the video started. One second after the response, the video started, and participants viewed in random order either a TI or an FT of either the flexible or the rigid actor, followed by a blank screen for 250 ms, then one of the probe pictures for 250 ms. Finally, the question “same position or not?” was written on the screen until the participants responded by pressing the key “o” for yes or “n” for no (cf. previous experiments). There was no time pressure or constraint. Trials were separated with a 1000 ms interval. Participants started the task with a training session including 24 FTs. Then, participants performed eight blocks of 60 trials interspersed with short breaks, for a total of 480 trials. In each block, there were 48 FTs (80%) that included the three probes in equal proportions intermixed with 12 TIs (20%) that included only forward and backward probes, in equal proportions, resulting in a total of 24 TIs per each combination of the two actors and two probes. The actors and the probes were randomized within blocks of trials. Additional details are listed in Table 1.

**Table 1.** Summary of the main parameters of Experiments 4 to 7.

|  | Experiment 4 | Experiment 5 | Experiments 6 & 7 |
| --- | --- | --- | --- |
| Software | Testable | Psychopy | Psychopy |
| Stimuli format | 60 Hz videos | 60 Hz videos | 1 frame by refresh rate (60 Hz) |
| **Probes** |  |  |  |
| FTs (n/block) | -5°/0°/+5° (16) | -10°/-5°/0°/+5°/+10° (*) | -10°/0°/+10° (5) |
| FLTs (n/block) | na | na | 0°/+10° (1) |
| TIs (n/block) | -5°/+5° (6) | -5°/+5° (6) | 0°/+10° (1) |
| CT (n/block) | na | -30° (2) | na |
| Number of blocks (n trials) | 8 (60) | 6 (60, +2 catches) | 10 (17, +2 fillers) |
| Actor order | Randomized within blocks | Randomized within blocks | Fixed by blocks |
| Response mode | Keyboard (o/n keys) | Keyboard (o/n keys) | Verbal (oui/non) |

*Note.* FTs = Familiarization trials; FLTs = Filler trials; TIs = Trials of interest; CT = Catch trials.
\*There were 16 repetitions of the identical probes and eight repetitions of each other probe.

After the task, participants were invited to try to guess the goal of the experiment. The experiment was controlled by the online testable.org interface (http://www.testable.org), with video-conference software to monitor the session and interact with the participants.

### Analyses and results

Two exclusion criteria were predetermined: (1) making more than 5% of errors in identifying the actor at the beginning of a trial and (2) correctly guessing the purpose of the experiment. No participant was excluded from the analyses.

The main objective of this experiment was to test the prediction that displacement should be different when participants see the flexible and the rigid actor performing an external rotation of 75 degrees (i.e., the TIs). To test this hypothesis, we ran a GLMM with the participants’ responses (yes/no) entered as the dependent variable and Probe (forward/backward) as well as Actor (flexible/rigid) as fixed within-subject factors. Participant was included as a random factor. In line with the prediction of the visual hypothesis, there was a significant interaction between Probe and Actor characterized by a larger skewness index (probability of “yes” response between forward minus that for backward probes) for the flexible actor (*M* = .23, *SE* = .03) than for the rigid actor (*M* = .01, *SE* = .03), *F*(1,1916) = 23.76, *p* < .001, *OR* = 2.60.

The results of additional analyses conducted to explore whether participants’ perception was characterized by forward displacement in this task indicated that this was the case for the flexible actor, *t*(1916) = 7.16, *p* < .001, but not for the rigid actor, *t*(1916) = 0.41, *p* = .68, providing additional evidence that perceptual extrapolation of body movements is actor-specific. This, and the fact that there was also a significantly larger probability of “yes” responses for the backward probes when participants observed the rigid actor (*M* = .59, *SE* = .04) than when they observed the flexible actor (*M* = .52, *SE* = .04), *t*(1916) = 2.08, *p* = .02, indicated that forward displacement was non-significant when participants observed the rigid actor because it was significantly smaller.

Importantly, the results of another exploratory analysis, conducted on a subset of the TIs for which participants’ responses were provided in less than one second (82%) were virtually identical: there was a larger probability difference of “yes” response between forward and backward probes for the flexible actor (M = .25, SE = .03) than for the rigid actor (M = .04, SE = .03), and the difference was significant, *F*(1,1553) = 17.68, *p* < .001, *OR* = 2.50. This suggests that this effect is unlikely to result from decision-related processes (see general discussion).

Note, finally, that we ran additional exploratory analyses of the data reported in this section, including analyses of variances (ANOVAs) and GLMMs that include the independent variables and their interaction as random factors. All the additional analyses led to the same conclusions (see detailed results on OSF).

**Figure 7.**
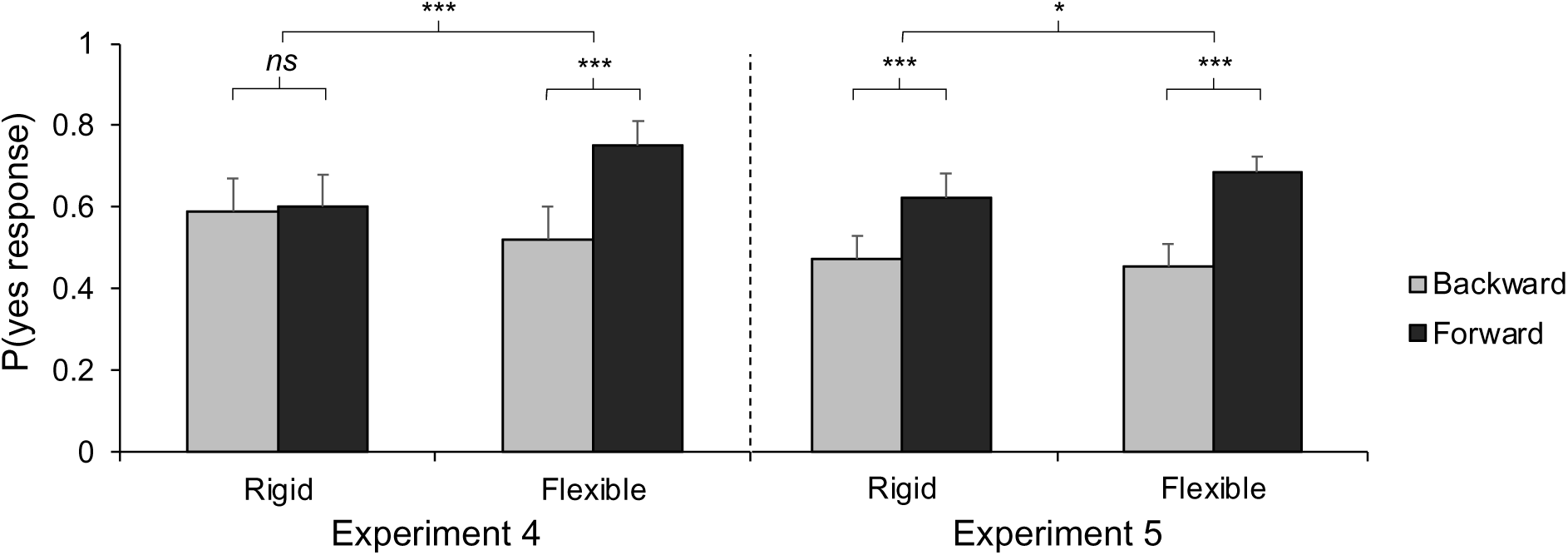
Results of Experiments 4 and 5. *Note*. Probability of a “yes” response as a function of Probe (forward/backward) and Actor (flexible/rigid) in Experiments 4 and 5. Error bars depict 2 standard errors of the mean. *ns*: non-significant, \**p* < .05, \*\*\**p* < .001

### Discussion

Three main findings emerged from this study: (1) the observation of the flexible actor was associated with a significantly larger number of “yes” responses for forward than for backward probes (skewness index); (2) the skewness index was significantly larger when participants observed the flexible than the rigid actor and (3) there was a significantly larger probability of “yes” responses for the backward probes when participants observed the rigid than the flexible actor.

These findings invite two main conclusions. The first finding implies that participants’ responses were influenced by forward displacement (FD) when observing the movement executed by the flexible actor. Indeed, it is unclear how this finding could be accounted for by the alternative hypotheses: (a) that there is no displacement, or (b) that there is significant backward displacement. Together, the two other findings imply that there was a significantly larger forward displacement evoked by the observation of the flexible than by the observation of the rigid actor. Again, the combination of findings (2) and (3) cannot be accounted by the two alternative hypotheses: (a) that the two actors evoke the same forward displacement, or (b) that in fact the rigid actor evokes a larger forward displacement than the flexible one.

This suggests that visual perception may be constrained by actor-specific expectations, in line with the predictions of the visual hypothesis of body movement perception. In contrast, these findings appear difficult to reconcile with the motor hypothesis of body movement perception. If extrapolation of others’ body movements were underpinned by an internal model of the observer’s own body (e.g., Knoblich et al., 2002; Wilson & Knoblich, 2005), there should be no difference between the displacement evoked by the two actors.

However, it remains unclear whether this influence results from participant’s visual experience with the two actors’ different body abilities, or from the explicit verbal information that was conveyed to the participant about the two actors’ flexibility. To address this issue, Experiment 5 was an attempt to replicate the results of Experiment 4 in a study in which participants learned about the actors’ different amplitudes of movement through visual experience only, without any explicit instructions.

## Experiment 5

### Method

#### Participants

A sample size of 40 participants (33 females, seven males, *M_age_* = 20.1, *SD* = 1.1) was determined a priori on the basis of a power analysis. This provided a power of at least 90% to detect an effect of the actor of a size of at least half that found in Experiment 4 (i.e., Cohen’s *d* = 0.5). All participants received course credit for their participation.

#### Stimuli and procedure

The important changes vs. Experiment 4 were the following: (1) during the instructions, participants received no information about the actors’ different movement amplitude. Instead of categorizing the actors as “flexible” or “rigid,” they were portrayed as the actors with either “black” or “blond” hair and the two actors were introduced to participants through a picture depicting their right arm tilted 20 degree of external rotation (i.e., the posture corresponding to the TIs); (2) in the FTs, the flexible actor performed an external rotation of 110 degrees (see Figure 6); (3) two catch trials were added to each block (always a FT for the flexible actor with a probe depicted his hand shifted 30° backward) and (4) the experiment was run in the laboratory. Other changes are listed in Table 1, which provides a summary of the main parameters of Experiments 4 to 7.

### Analyses and results

No participant was excluded based on our a priori exclusion criteria. Like in Experiment 4, we found a significant interaction between Probe and Actor characterized by a larger skewness index for the flexible actor (*M* = .24, *SE* = .03) than for the rigid actor (*M* = .16, *SE* = .03), *F*(1,2876) = 4.47, *p* = .01, *OR* = 1.40.

Exploratory analyses further indicated that participants’ perception was characterized by a forward displacement (i.e., a significantly larger number of “yes” responses for forward than for backward probes) for both the flexible actor, *t*(2876) = 9.25, *<u>p</u>*< .001, and the rigid actor, *t*(2876) = 6.31, *p* < .001. And like in Experiment 4, there was no indication that the smaller skewness index associated with the rigid actor may have emerged in the context of a larger forward displacement, as it was not associated with a smaller probability of “yes” responses for the backward probes, *t*(2876) = 0.33, *p* = .75.

Like in Experiment 4, the results of an exploratory analysis conducted on a subset of the TIs for which participants’ responses were provided in less than one second (78%) were also very similar: there was a larger probability difference of “yes” response between forward and backward probes for the flexible actor (*M* = .26, *SE* = .03) than for the rigid actor (*M* = .17, *SE* = .03), and the difference was significant, *F*(1,2244) = 4.21, *p* = .02, *OR* = 1.45. This suggests that this effect is unlikely to result from decision-related processes (see general discussion).

Note, finally, that we ran additional exploratory analyses of the data reported in this section, including analyses of variances (ANOVAs) and GLMMs that include the independent variables and their interaction as random factors. All the additional analyses led to the same conclusions (see the detailed results on OSF).

**Figure 8.**
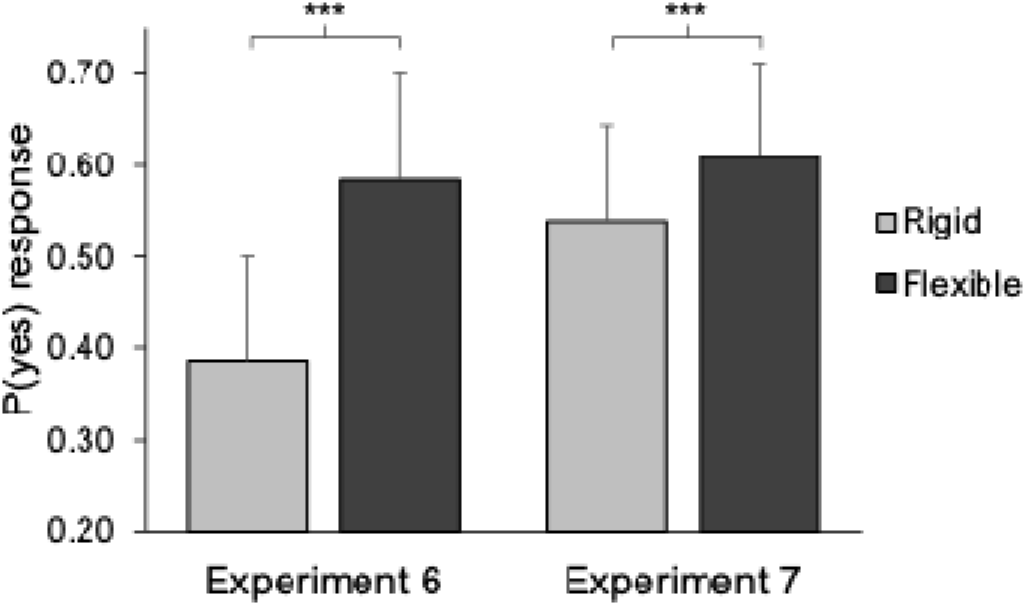
Results of Experiments 6 and 7. *Note*. The probability of a yes response for forward probes, illustrated as a function of Actor across both experiments. Error bars depict 2 standard errors of the mean. \*\*\**p* < .001

### Discussion

The results of this experiment were virtually identical to those of Experiment 4. There was a significantly larger number of “yes” responses for forward than for backward probes for the flexible actor (skewness index). The skewness index associated with the flexible actor was significantly larger than that associated with the rigid actor. And there was there was a significantly larger probability of “yes” responses for the backward probes when participants observed the rigid than the flexible actor. Like in Experiment 4, these results provide evidence that participants’ responses were influenced by a significant and significantly larger forward displacement (FD) when observing the movement executed by the flexible than by the rigid actor. These results, in line with the predictions of the visual hypothesis, suggest that perceptual extrapolation of body movements is shaped by knowledge of others’ unique biomechanical constraints.

It remained unclear, however, whether participants were influenced by visually acquired models of the two actors’ biomechanical constraints (i.e., what they can do) or, rather, by expectations based on their experience of how the two actors move in the previous trials of the experiments (i.e., what they do in the task). In these experiments, one actor often moved (i.e., in the familiarization trials) past the endpoint of the TIs while the other did not. The goal of Experiments 6-7 was to disentangle these two interpretations. To do so, internal rotations of the shoulder were used as FTs (instead of external rotations in Experiments 4 and 5). This way, participants saw the exact same external rotations (in the TIs) for both actors and a differential effect could not be attributed to task-related repeated patterns.

## Experiment 6

### Method

#### Participants

A sample size of 25 participants (15 females, 10 males, *M* = 23.1, *SD* = 4.7) was determined a priori on the basis of a power analysis. This provided a power of 90% to detect an effect of the actor of a size of Cohen’s *d* of 0.6, which was determined based on the result of a pilot study involving 11 other participants. To reach this number, we tested 37 participants, of whom 12 were excluded based on a priori exclusion criteria. All of them received course credit for their participation.

#### Material and procedure

This experiment was very similar to Experiment 5 (cf. Table 1). The important changes were the following: (1) in the FTs, actors performed internal rather than external rotations of the right shoulder (cf. Figure 6); (2) filler trials depicting external rotations of the shoulder within the range of motion of TIs (60° rotations) were added to ensure that participants could not predict the end-point of external rotations; (3) TIs were followed by either forward or identical probes. The use of identical probes ensured that external rotations were followed by a similar number of correct “yes” and “no” answers, and discarding backward probes allowed us to shorten the task without affecting our ability to test a differential prediction of the visual and motor hypotheses (i.e., the presence, or not, of a difference between the two actors); (4) actors were blocked; (5) the participants were familiarized explicitly with the two actors’ maximal amplitude rotation in the same way as in Experiment 4; (6) directly after this familiarization (and before starting the localization task), participants were asked to reproduce the maximal amplitude of each actor (maximal amplitude judgment task; MAJ task). Participants were shown one actor in the starting position of TIs on a picture and were then required to press the right and left arrows of the keyboard to navigate through frames depicting the actors’ shoulder rotated toward or away from the body, from 30° clockwise (starting position in TIs) to 70° counterclockwise (10° beyond the maximal amplitude of the flexible actor) by steps of 5° and press the space bar to validate the position that matched the maximal amplitude of that actor. Then, they did the same task for the second actor. Any error at this point led the participant to a new attempt, with a maximum of four attempts; (7) during the localization task, participants gave their response orally (yes/no) and then pressed the space bar to launch the next trial; (8) participants were not asked to identify the actor before trials.

### Analysis and results

Twelve participants who failed the MAJ task after four attempts were excluded (a priori exclusion criterion) and replaced. No participant was able to guess the purpose of the experiment.

Data were analyzed by a generalized linear mixed model (GLMM) in which the responses (yes/no) were encoded as the dependent variable. As there were no backward probes, we analyzed the responses to the forward probes as a function of the Actor (flexible/rigid), entered as a within-subject fixed factor. Participant was included as a random factor. In line with the visual hypothesis, the analysis revealed a main effect of the actor, characterized by a larger probability to respond yes to a forward probe for the flexible actor (*M* = .58, *SE* = .06) than for the rigid actor (*M* = .39, *SE* = .06), *F*(1,498) = 16.78, *p* < .001, *OR* = 2.24. These results were replicated with analyses of variances (ANOVAs) and GLMMs including the independent variables and their interaction as random factors (see on OSF).

### Discussion

The results of Experiment 6 confirmed that actor-specific perceptual extrapolation may be based on internal models of these actors’ biomechanical constraints rather than on task-related expectations. The goal of Experiment 7 was to replicate the results of Experiment 6 in a design in which participants learned the two actors’ different amplitude of movement only incidentally, through their visual exposure to the FTs, without any explicit instruction.

## Experiment 7

### Method

#### Participants

A sample size of 36 participants (29 females, seven males, *M* = 21.7, *SD* = 3.6) was determined a priori on the basis of a power analysis. This provided a power of 90% to detect an effect of the actor of a size of Cohen’s *d* of 0.5 (i.e. half the size of the effect size detected in Experiment 6). To reach that number, we tested 45 participants, of which nine were excluded based on criteria established before the study (see results section). All of them received course credit for their participation.

#### Material and procedure

The changes from Experiment 6 to the present experiment were the following: (1) no indication regarding the actors’ maximal amplitude was given to participants before the experiment. During the instruction, participants were presented with a picture displaying the two actors in the same posture (the endpoint position of the TIs); (2) participants were only invited to perform the maximal amplitude judgment task for both actors at the end of the localization experiment (instead of before in Experiment 6).

### Analysis and results

Among the 45 students who participated in the experiment, nine were excluded before any data analysis because they reported a larger amplitude of movement for the rigid than the flexible actor in the MAJ task. No participant was able to guess accurately the purpose of the experiment. We ran the same GLMM analysis as in Experiment 6, in which the responses of the TIs to the forward probe (yes/no) were encoded as the dependent variable, and the Actor (flexible/rigid) was entered as a within-subject fixed factor. Participant was included as a random factor. The analysis revealed a main effect of the actor, characterized by a larger probability to respond yes to a forward probe for the flexible actor (*M* = .61, *SE* = .05) than for the rigid actor (*M* = .54, *SE* = .05), *F*(1,718) = 3.3, *p* = .03, *OR* = 1.35. Again, these results were replicated with analyses of variances (ANOVAs) and GLMMs including the independent variables and their interaction as random factors (see on OSF).

### Discussion

In this experiment, as in Experiment 6, we observed a significantly higher number of “yes” responses for forward probes when participants viewed the flexible actor compared to the rigid actor. This indicates that visual perception may be constrained by actor-specific expectations, in line with the predictions of the visual hypothesis of body movement perception. In contrast, these findings appear impossible to account for by the current versions of the motor hypothesis of body movement perception, which suggests that the biomechanical bias in body movement perception is underpinned by an internal model of the observer’s own body (e.g., Knoblich et al., 2002; Wilson & Knoblich, 2005). If the motor hypothesis were correct, we would expect no difference in displacement between the two actors.

These results therefore corroborate and extend those of Experiments 4 and 5. The results of Experiments 6 and 7 cannot be explained as a consequence of mere expectations resulting from the fact that one actor was often seen executing external rotations of a larger amplitude than the other one. Rather, they suggest that incidental visual learning of a model specifying the actors’ different ranges of possible shoulder movement (i.e., biomechanical constraints) may shape the perception of these actors’ body movements.

Despite the theoretically important differences detected between the percepts evoked by the observation of the same movement executed by the two different actors, the probe task used in Experiments 6 and 7—with only a single forward probe—did not allow us to determine whether the stimuli elicited significant forward displacement (limitation 1).

This design also prevented us from ruling out the possibility that the higher number of “yes” responses for the forward probe associated with the flexible actor might be explained by a larger forward displacement for the rigid actor (limitation 2). Such a displacement could have extended beyond the location tested by our single probe, potentially reducing the number of errors for the rigid actor.

When designing Experiments 6 and 7, we considered these limitations minor and not compelling enough to justify the inclusion of a backward probe. First, and most importantly, the inclusion of a backward probe was not necessary to test the critical differential prediction of the visual and motor theories tested in Experiments 4-7, which is centered on whether or not the perception of the two actors would evoke different responses, rather than on whether the forward displacement itself was significant (limitation 1) or on confirming that the flexible actor was indeed associated with the largest forward displacement (limitation 2). Second, the results of Experiments 3-5 already provided clear and robust evidence that the perception of the flexible actor does indeed evoke significant forward displacement. Third, the hypothesis that the rigid actor might exhibit a larger forward displacement in Experiments 6 and 7 (limitation 2) seemed improbable. Indeed, such effect would only contradict the results of Experiments 3-5, where the skewness index was consistently larger for the flexible actor, and the percentage of “yes” responses to the backward probes was consistently higher for the rigid actor. Moreover, this outcome would conflict with the predictions of both motor and visual theories of body movement perception and with any theoretical framework we considered.

Nevertheless, to address these issues, Experiment 8 employed a pointing task to assess participants’ perception of the final position of the two actors’ hand. This approach allowed for a conceptual replication of the findings reported in Experiments 4-7 and provided a more direct and fine-grained measurement of participants’ perceived final positions of the two actors’ hands.

## Experiment 8

### Method

#### Participants

We tested 58 participants (53 females, five males, *M_age_* = 20.9, *SD* = 4.2), which provided us a power of at least 0.8 to detect a Cohen’s D of at least 0.3 for the actor effect (i.e., a small effect size). All participants received course credit for their participation.

#### Material and procedure

Stimuli were the same four video-clips as in Experiment 6 and 7 (see Figure 6), showing the two actors performing either an internal (FT) or an external (TI) rotation of the shoulder. In each trial of the experiment, the participant saw one of the four video clips followed by a small red dot (5×5 pixels) on the center of an otherwise blank screen. They were asked to carefully look at the video-clip, then to use the computer mouse to move the small red dot (mouse cursor) as exactly as possible to where the tip of the middle finger of the actor was at the end of the video-clip, and then to press a button on the computer mouse to validate their response. There were 10 blocks of 34 trials interspersed with short breaks, for a total of 340 trials. Blocks were composed of 30 FTs and four TIs (see Figure 6) presented in random order. The actors were blocked, and the order of the blocs was counterbalanced across participants. At the end of the experiment, participants were invited to try to guess the goal of the experiment.

### Analysis and results

No participant was able to guess correctly the goal of the experiment. Only the TI trials (external rotations) were analyzed. Then, data analysis proceeded in four steps.

First, TIs in which the Euclidean distance between the recorded and actual location of the actor’s middle finger deviated by more than 2.5 median absolute deviations (Leys et al., 2013) from each participant’s median distance in each condition (FLEXIBLE/RIGID) were excluded from the analysis (4.1% of the trials).

Second, for each remaining trial, we identified the point along the (past or future) trajectory of the actor’s middle finger that was the closest to the coordinates of each of the participant’s response (closest point, CP). To identify the closest point (CP), we first applied a polynomial function to interpolate points in the plan through which the hand had passed (before the movement stopped) or would have passed (if the movement had continued), based on the actor’s hand position across the frames of the videos. Then, we identified the coordinates of the closest point along this trajectory to each of the participant’s responses. Finally, we calculated the arc length (in degrees of visual angle) between each CP and the true final location of the hand and added a sign such that a positive value indicated a shift forward along the motion trajectory, and a negative value indicated a shift backward along the motion trajectory. Hence, a trial associated with an arc of 2 degrees indicated that, in that trial, the participant had located the final position of the middle finger of the actor closer to where the middle finger would have been if it had continued to move for 2 degrees along its trajectory after the video disappeared than to any other point along the past or future position of the finger.

Third, we ran a generalized linear mixed model (GLMM) including the signed arc length as the dependent variable, the Actor (flexible/rigid) as a within-subject fixed factor, and Participant as a random factor in order to test whether there was a difference in the location of the last position of the middle finger of the two actors, as predicted by the visual hypothesis. This was the case. There was a significantly larger positive arc length for the flexible (*M* = 0.25, *SE* = 0.05) than for the rigid actor (*M* = 0.16, *SE* = 0.05), *F*(1,2223) = 32.76, *p* < .001.

Finally, we ran one-sample t-tests and found that the average signed arc lengths were significantly larger than zero for both the flexible, *t*(1106) = 15.67, *p* < .001, and the rigid actor, *t*(1117) = 9.35, *p* < .001. In other words, on average, the participants perceived the final position of both actors’ hand closer to where it would have been if it had briefly continued to move along its initial trajectory after the video-clip ended than to any other points – a forward displacement.

All these results were replicated with analyses of variances (ANOVAs) and GLMMs including the independent variables and their interaction as random factors (see on OSF).

### Discussion

These results therefore corroborate the findings and conclusions from Experiments 4—7. In line with the prediction of the visual hypothesis, there was once again a significant difference between the perceived final location of the two actors’ middle finger (like in Experiments 4-7) and this difference was characterized by a significantly larger forward displacement elicited by the observation of the flexible than the rigid actor (like in Experiment 4-5).

## General Discussion

Visual perception of human body movements is constrained by knowledge of the body biomechanics (Shiffrar & Freyd, 1990; Vandenberghe & Vannuscorps, 2023; Wilson et al., 2010). A critical issue concerns whether such effect originates from a model of one’s own body, acquired through motor experience and proprioceptive feedback, or from models of others’ bodies, learned through visuo-perceptual experience. Here, we have reported the results of eight experiments conducted to test several differential predictions of these two hypotheses. All these results were compatible with the predictions of the “visual hypothesis” but incompatible with the predictions of the “motor hypothesis”. First, we have reported a typical influence of knowledge of the upper limb biomechanics on apparent movement perception (Experiment 1) and perceptual extrapolation (Experiment 2) in individuals completely deprived of upper-limb sensorimotor experience from birth. Second, we have reported clear evidence that observers extrapolate significantly less the perception of a movement that would have been difficult to continue for an average actor, even if the same movement would have been easy to continue for the observers themselves (Experiment 3). Third, we have reported that body movement perception is shaped by knowledge of how different persons can move, learned either through explicit instructions (Experiments 4 and 6) and/or incidentally through mere visual experience (Experiments 5, 7, and 8).

These findings complement those of two previous studies that had documented an influence of knowledge of the typical upper limb biomechanics in apparent motion and localization tasks in individuals born without hands (Vannuscorps & Caramazza, 2016a, 2016b). In these studies, however, the effect of the biomechanical constraints could have emerged from these individuals’ sensorimotor experience with their lower limbs (see discussion of Experiments 1 and 2). In addition, the differential predictions of the visual and motor hypotheses remained to be tested in typically developed participants. The individuals born without upper limbs tested in Experiments 1 and 2 were also deprived of lower limbs, and the results of Experiments 3-8 provided strong support to the (visual) hypothesis that perceptual extrapolation depends on the observer’s knowledge of the actors’ biomechanics rather than on his/her own body ability.

These conclusions are in line with those of a large set of previous studies indicating that observers anticipate the future posture of an observed action (Imanaka, Sugi, & Nakamoto, 2023; Verfaillie & Daems, 2002; Wilson et al., 2010; Zucchini, Borzelli, & Casile, 2023) if it is predictable (Munger, 2015) and that expectations influence displacement (e.g., Finke et al.,1986). For instance, Hudson and his colleagues (2018) showed that when the few first frames of a hand reaching for an object are depicted, the presence or absence of an obstacle further along the path affects the reported final position of the hand. When participants were instructed to indicate where the hand disappeared on a touchscreen, their responses were consistently shifted toward the most efficient trajectory. Specifically, the vertical shift was higher when the obstacle was present, as the hand aimed to avoid it, compared to when the obstacle was absent, as the hand aimed to reach straight for the object.

The results of the eight experiments reported here also provide new insight on the mode of acquisition, types, and nature of the models of the body biomechanical constraints that may influence perception: they suggest that our visual system learns incidentally (Experiments 5, 7 and 8) and uses automatically both generic and actor-specific models of the human body biomechanics to guide perception. Hence, in Experiments 1-3, like in previous studies, participants were not given any information or experience about the observed actor’s flexibility. However, the end position of the actor’s arm was set at 20 degrees of external rotation of the shoulder, which corresponds to the average maximal amplitude of external rotation of the shoulder for young adults (see the results of Experiment 3; see also Gill et al., 2020; Hill et al., 2010). Thus, that participants showed a reduced forward displacement for external rotations of the arm in these experiments suggests that perception may rely on a generic model of how typical young adults can and cannot move – that participants implicitly “knew” that it would be unlikely that the depicted actor would be able to continue to move his arm further along the same trajectory.

The results in Experiments 4-8 additionally indicate that observers may also rely on more specialized “actor-specific” models when relevant information is available. Previous studies have demonstrated that we can recognize people that we know based on “how they move” (Cutting & Kozlowski, 1977; Loula et al., 2005) and that we are able to identify and discriminate people by their gait after very little observation of their movements (Jacobs et al., 2004; Troje et al., 2005). Here, we show that our visual system encodes and uses not only information about how others typically move but also about the range of movements they can perform.

Hence, in addition to contributing to the debate about the motor vs. visual nature of the effect of biomechanical plausibility on perception, the findings reported herein open new research avenues. The existence of “generic” models begs several new questions. The actor depicted in Experiments 1-3 was a young adult and the movement depicted corresponded to the average maximal range of external rotation of young adults. Yet, the maximal range of motion is influenced by many factors such as age, gender, physical activity, fitness, and injury history.

Although knowledge of young male adults’ average maximal range of movement is likely a key to generating accurate visual extrapolations when observing actors who belong to that category, relying on this knowledge would probably not be optimal when observing body motion from other categories of persons, such as children and older adults, who are typically more and less flexible, respectively. Does it mean that we learn, encode, and use models of the body biomechanics of different “categories of people”? If so, what determines these “categories”? And how are the models of the biomechanical constraints of a category of person encoded and defined? One possibility that we can think of is that generic models represent merely the average range of motion of individuals from a given category that an observer has previously encountered. Another possibility is that generic models represent a probability function based on the distribution of the maximal range of movement of all the familiar persons from a category. Similar questions may be raised for actor-specific models. Another question relates to the neural substrate of these models and their influence on perception. Previous research using functional magnetic resonance imaging (fMRI) points toward the Extrastriate Body Area (EBA) as a candidate for housing generic models. Although the precise location of EBA varies slightly between individuals, it is generally found in the lateral occipitotemporal cortex, near the junction of the occipital and temporal lobes (Downing et al., 2001). EBA responds to pictures of arms, hands, and torsos significantly more than to other types of visual stimuli such as man-made objects, animals, and faces (Downing et al., 2006; Downing et al., 2001; Peelen et al., 2006) and it responds differently to pictures of natural vs. strongly distorted and unfamiliar finger postures (i.e., fingers bent backward, Avikainen et al., 2003) and to video-clips of natural vs. biomechanically impossible finger movements (i.e., an abduction of approx. 90 degrees of the pinkie; Costantini et al., 2005). The results of one recent study indicate that EBA may also encode models of actor-specific body biomechanics (Bellot et al., submitted). In that study, we first presented participants with information about the maximum amplitude of arm and leg movements exhibited by both a “rigid” and a “flexible” actor. Then, we assessed the BOLD response in the same participants when they viewed video clips portraying these actors performing three types of movements: “small” and attainable by both actors, “large” and beyond the capabilities of both actors or “intermediate” and attainable only by the “flexible” actor. In line with the hypothesis that EBA encodes models of actor-specific biomechanics, Multivoxel pattern analysis of the fMRI data (MVPA) revealed that the EBA categorized intermediate movements as either possible or impossible based on the actors’ individual abilities.

More research is also required to develop and test hypotheses about the precise cognitive origin of the effect of knowledge of the body biomechanics in the apparent motion and localization tasks. In all the experiments reported above, we ensured that participants were neither aware of the goal of the study nor able to correctly guess it. This reduces the possibility that our results reflect mere demand characteristics (Orne, 1962).

At first sight, it may be tempting to suggest that the influence of knowledge of the body biomechanics on the perceived path of apparent motion (Experiment 1) could result from decision-related processes. The effect would emerge because, when faced with the need to provide a response, participants reason about the biomechanical plausibility of the movements depicted on the response screen (Figure 1A) and decide to select the biomechanically plausible movement. Several aspects of the methods and results of our experiment make this hypothesis highly unlikely, however. First of all, decision-related effects are typically reported in tasks in which participants seek a correct answer among several conflicting options. In our experiment, however, participants were not forced to choose between the two motion paths. They had the opportunity to respond that they did not see any motion or not one of the two motion paths depicted on the response screen. And, importantly, before the experiment they were familiarized with the fact that perception of the same stimulus can differ from time to time and from person to person and told to “relax and tell us what you see” because “there is no correct or incorrect response”. In our opinion, it is therefore unlikely that participants engaged in any sort of reasoning or strategy when providing their responses. Indeed, no participant reported doing so when interviewed at the end of the experiment.

A more likely hypothesis, in our opinion, is that knowledge of the body biomechanics influences the perceived path of motion because this information is used as high-level top-down information that shapes the trajectory of the illusory path through feedback connections to earlier visual areas. Indeed, apparent motion is largely assumed to be the result of a constructive process by which the visual system “fills in” missing information. Accordingly, apparent motion elicits retinotopic activity in V1 that reflects the subjective “illusory” path of motion perceived by the participants (e.g., Akselrod et al., 2014; Muckli et al., 2005; Shen et al., 2020) and impairs the detection of visual targets located along the illusory path (Hidaka et al., 2011; Liu & Lourenco., 2021; Shen et al., 2020). In other words, the object is literally processed as if it had really appeared at intermediate locations along the path of motion, where it was never displayed. Although this hypothesis remains to be tested empirically, its plausibility is also supported by the results of studies indicating that V1 activity during apparent motion is driven by higher-order parts of the visual system, in particular the human middle temporal complex (hMT+; Sterzer et al., 2006; Vetter et al., 2015), which overlaps partly and is strongly connected to EBA (e.g., Ferri et al., 2013). If this hypothesis is correct, apparent motion of the body should impair the detection of targets placed on the trajectory of a biomechanically plausible path of motion, rather than on the shortest path of motion, when the two differ, and transcranial magnetic stimulation of EBA should hamper the biomechanical bias.

The origin of the effect of knowledge of the body biomechanics in the localization task is less clear and probably multi-determined (Hubbard, 2010; Kerzel, 2005). Given the difficulty of the task, it is possible that under uncertainty participants strategically call into play various sources of information to guide their decisions, including information about a person’s body ability. For instance, when presented with a “forward” probe picture, under uncertainty participants may eventually decide to respond more often “yes” (“this was the position of the actor’s hand at the end of the video”) if they know that the actor would have been able to reach that position than if they think that the actor would not. However, several arguments suggest that the effects that we report are unlikely to be reducible to an effect of knowledge and expectations on strategic, conscious decision-related processes.

First of all, there is clear evidence that displacement does not result from, and is not easily influenced by, decision-related processes. The most convincing support for this conclusion comes from studies showing that forward displacement survives participants’ conscious, strategic attempts to control it. Ruppel and colleagues (2009) showed that displacement resists even feedback specifying the direction of errors, for instance (see also Finke & Freyd, 1985; Freyd, 1987). In another study, Courtney and Hubbard (2008) tested displacement induced by the implied motion of a target in three groups of participants. Participants from the first group were not given any particular information. Participants from the second group were informed beforehand that the task is known to elicit displacement. In the last group, participants were additionally instructed to try to compensate for displacement in their responses. The logic of the study was rather straightforward: if displacement results from decision-related processes, then, it should disappear (or even reverse) when participants are informed about it or asked to counteract it. In contrast to this hypothesis, instructions reduced but did not eliminate displacement in any of the groups.

In addition, there is also some evidence that displacement is influenced by participants’ implicit rather than explicit knowledge (Freyd & Jones, 1994; Kozhevnikov & Hegarty, 2001). In a particularly interesting experiment, Kozhevnikov and Hegarty (2001) compared the displacement elicited by ascending stimuli of different mass in participants who were either naïve or experts of Newtonian physics. All the participants from the expert group stated beforehand that heavier objects should ascend more quickly than lighter objects, in line with Newtonian physics. The participants from the naïve group, however, all believed that lighter objects would ascend more quickly than heavier objects. Despite this striking difference in explicit knowledge, the heavy ascending target produced significantly less forward displacement than the light one in both groups, which did not differ by any measure. That implicit rather than explicit knowledge or expectations influence displacement seems difficult to reconcile with the hypothesis that this influence emerges from a conscious, strategic attempt of participants to reason about the most likely correct response to provide in this task.

In addition to these theoretical considerations, two aspects of our own findings seem to challenge that hypothesis. First of all, as discussed above, at the end of the experimental session of Experiments 5 and 7, when asked whether they used any particular strategies to help them solve the task, none of the participants declared having used information about the actors’ ability. Second, in Experiments 4-5, participants provided the majority of their responses (>78%) in less than one second and exploratory analyses of these responses only led to the same conclusion than the analysis of the whole set. This seems difficult to reconcile with the idea that the effect emerges due to strategic, conscious decision-related processes, which involves a series of time-consuming steps that arguably take more than 1000 ms: (1) looking at the probe picture and realizing that one is not sure whether the probe depicts the hand where it was at the end of the video-clip or not, (2) reason about whether the position would have been possible or not for that particular actor, (3) and use this information to guide their response.

It is possible, however, that knowledge alters the representation of the correctly perceived final position of the actors’ moving hand while it is maintained in memory during the interstimulus interval. Indeed, displacement is often assumed to result from distortions of representations held in memory (Freyd & Finke, 1984). In line with this hypothesis, displacement has been documented with static pictures. For instance, in a study by Freyd (1983), participants were shown one photograph of an action like “jumping from a wall” or “a seagull flying” for 250 ms, an interstimulus interval of 250 ms, and then either the same picture or a picture of the same action taken a few milliseconds (between 55 and 500 ms) before or after the first one. Participants had to decide as rapidly as possible whether a second picture was “same as” or “different from” the first. Subjects took longer to indicate that the second picture was different when it depicted the action photographed slightly after the first one than when it depicted it slightly before the first one (see also Freyd et al., 1988; Futterweit & Beilin, 1994, for similar results on error rates). Freyd (1983) concluded from these findings that when subjects perceived the first photograph, they mentally “continued them” forward in time in the direction of the implied motion. As a result, memory for the first picture is more similar to that of the picture depicting it displaced forward in time. It is therefore possible that knowledge of the body biomechanics shapes the mental representation of the final position of the actors’ hand in memory.

We see at least two other possible, non-mutually exclusive, ways by which knowledge of generic and actor-specific body biomechanics may influence participants’ responses in the localization task. The first is that the effect that we report emerges from an influence of knowledge on oculomotor behavior. The logic of this account is as follows. When observers are asked to localize the final position of the actor’s hand, they may try to follow the hand with their eyes. Obviously, eye movements may not be stopped instantaneously when a stimulus disappears. In fact, the eye continues to move for up to 300 ms after the target disappearance (Kerzel et al., 2001; Mitrani & Dimitrov, 1978; Stork & Müsseler, 2004). Because the image of the target may persist on the retina for about 60 ms, the distance covered by the eyes during this time may result in a localization error (Kerzel, 2000). In turn, the effects that we report could be accounted for under the assumption that knowledge of the actors’ maximal range of motion triggers a signal halting the oculomotor pursuit as the hand approaches the expected final position, decreasing the size of the overshoot — and thus of the size of displacement. In line with this hypothesis, smooth pursuit stopping is known to be strongly influenced by active prediction of the upcoming end of the target (Missal & Heinen, 2017) and it has been shown that when observers track a smooth motion visual target, the size of the mislocalization of the endpoint of that target is tightly coupled with the size of the oculomotor overshoot (see Kerzel, 2000; Stork & Müsseler, 2004).

A second possibility is that knowledge influences the outcome of perceptual processing per se. Top-down expectations may guide the online extrapolation of the real position of others’ body parts to compensate for the lag imposed by neural transmission and processing time (i.e., an explanation similar to that evoked above to explain the effect of knowledge of the body biomechanics on apparent motion). This hypothesis is supported by evidence that expectations do indeed induce anticipatory activity in the visual system (Blom et al., 2020; Ekman et al., 2017; 2023; Johnson et al., 2023). For example, in an fMRI study by Ekman and colleagues (2017), participants first saw for a 4-min period a dot moving rapidly (40 deg/sec) from the left to the right side of a computer screen. Then, after this familiarization period, they occasionally saw only the starting or end point of the sequence. As expected, the analysis of the fMRI data indicated that physically presenting the moving dot triggered sequential activity at the corresponding retinotopic locations in the primary visual cortex (V1) as well as activity in higher-level visual areas encoding motion such as motion-sensitive area hMT+. More interestingly, seeing only the starting point of the sequence (but not the end point) elicited a very similar pattern of activity in both V1 and hMT+, suggesting that predictions may lead to anticipatory activity in the path of future stimulus events. Intriguingly, a large proportion of participants also reported seeing something similar to a stripe after being exposed to the starting point of the sequence (Ekman et al., 2017), suggesting that such predictive activity could lead to the (mis)perception of something that never happened. Further support for this hypothesis comes also from the finding that interfering with feedback signals from higher-level visual areas to V1, by applying TMS on hMT (Maus et al., 2013; Senior et al., 2002) or by using dynamic visual noise masking (Hudson et al., 2018), severely decreases forward displacement and its modulation by expectations.

In conclusion, apparent motion and perceptual extrapolation of the most likely future trajectory of a moving object are two perceptual phenomena that are widely assumed to reflect attempts by the visual system to overcome the intermittent and delayed nature of visual information (Blom et al., 2020; Nijhawan, 1994). Here, we show that these two phenomena are modulated by generic and actor-specific body biomechanics knowledge.

## Constraints on Generality

Together, the eight experiments reported here indicate that knowledge of body biomechanics influences body movement perception in various experimental designs (including apparent motion and localization tasks) and among participants with different physical abilities (more or less flexible), body schemas (typical, born without limbs with and without phantoms), and ages (ranging from 20 to 60 years old). We interpret this effect as the manifestation of a strategy by which the visual system relies on a universal physical principle (a limb cannot move past its joint limit) to cope with a universal problem (i.e., delays in the processing of visual information). These findings, along with their interpretation, lead us to hypothesize that this effect is likely to generalize to the population at large. In line with this hypothesis, previous research has documented the influence of other universal physical principles (e.g., gravitation, friction) on forward displacement in groups of young adults from different continents (Europe: De Sá Teixeira et al., 2017; North America: Hubbard, 1997; Asia: Nagai et al., 2002), as well as in individuals diagnosed with schizophrenia (De Sá Teixeira et al., 2013).

Nonetheless, it is important to acknowledge that the participants tested herein, and in previous studies cited in support of our hypothesis, represent only a specific demographic within the population, mostly educated 20 to 30-year-old students. Therefore, the generalizability of our conclusions to individuals outside this demographic remains to be verified. In addition, and to avoid misunderstanding, it may be useful to clarify that our hypothesis bears on the universality of the influence of knowledge of the body biomechanics on body movement perception. It predicts that we should be able to find traces of this influence across the population. Nonetheless, we expect the influence of knowledge of the body biomechanics to be modulated by a series of general factors known to influence forward displacement such as age (Hubbard, 2005) and attentional abilities (Hayes & Freyd, 2002). In addition, we would expect that the effect be modulated by individuals’ idiosyncratic knowledge and expectations, based on their individual experience and beliefs regarding body abilities and the observed body movement. We would predict that dancers and gymnasts who are often exposed to larger-than-average lower-limb movements, for instance, will likely overestimate the typical biomechanical limits of lower limb movements and, therefore, extrapolate the position of a moving lower limb significantly more than physiotherapists who, in contrast, may have more accurate models of generic and actor-specific biomechanics.

A last issue concerns whether our conclusion generalizes to the perception of other types of objects. The results of previous studies on whether displacement may be influenced by knowledge of the typical motion of other types of objects are either unclear (Reed & Vinson, 1996; Vinson & Reed, 2002) or at odds with this suggestion (Nagai and Yagi, 2001; see Vandenberghe & Vannuscorps, 2023 for discussion). In contrast, previous research has demonstrated that judgments of hand and body postures may also be influenced by knowledge of the body biomechanical limits (Parsons, 1987; Verfaillie & Daems, 1999, 2020), including in individuals congenitally deprived of the relevant limbs (Vannuscorps et al., 2013; Vannuscorps & Caramazza, 2015). Hence, we would expect perception of body postures to be influenced by knowledge of different actors’ body biomechanics, but this remain to be verified.

